# Temporal transcriptome profiling of *Microchloropsis gaditana* CCMP526 under hyper-saline conditions

**DOI:** 10.1101/2020.06.07.139238

**Authors:** Anbarasu Karthikaichamy, John Beardall, Ross Coppel, Santosh Noronha, Sanjeeva Srivastava, Dieter Bulach

**Affiliations:** IITB-Monash Research Academy, IIT Bombay, Mumbai 400076, India; School of Biological Sciences, Monash University, Clayton 3800, Victoria, Australia; Department of Microbiology, Monash University, Clayton 3800, Victoria, Australia; Department of Chemical Engineering, IIT Bombay, Mumbai 400076, India; Department of Biosciences and Bioengineering, IIT Bombay, Mumbai 400076, India; Medicine, Dentistry and Health Sciences, University of Melbourne, Melbourne 3010, Australia

## Abstract

Microalgae can tolerate a wide range of environmental conditions and have been exploited for their lipid and carbohydrate accumulating properties. The utility of this process could be further enhanced through understanding the critical gene regulatory networks that govern the acclimatization process. Advancements in systems biology and sequencing tools now enable us to obtain a genome-wide overview of gene expression under particular conditions of interest. Under salinity stress, *Microchloropsis gaditana* CCMP526, a commercially important alga has been previously reported to accumulate carbohydrate and lipid. To understand the mechanism of acclimatization, here we report a temporal transcriptomic analysis of *M. gaditana* under two different salinity levels (55 and 100 PSU). The short term (0, 1 and 6 h) and long term (24 and 72 h) responses of the salt-induced transcript pool were used to identify salinity-inducible genes using correspondence analysis. The transcript abundance of genes involved in triacylglycerol biosynthesis, membrane lipid modification, carbon assimilation and shunting, and osmolyte biosynthesis indicated that *M. gaditana* employs a two-stage acclimatization strategy during hypersaline conditions.

## Introduction

Increasing global energy demand driven by population growth and extensive dependence on fossil fuels, now recognized as a limited resource, has opened doors for renewable and sustainable energy sources (Georgianna & Mayfield, 2012). Researchers and several industries have embraced microalgae as an environmentally-friendly alternative fuel source to fossil fuels due to their high maximum growth rate, CO_2_ fixing capabilities and ability to tolerate environmental stress (Chisti, 2007; Moody et al., 2014; Stephens et al., 2010). Moreover, microalgae can accumulate high amounts of lipid under stress condition, and this phenomenon has been extensively researched in promising bio-fuel candidates under various stress conditions (Corteggiani Carpinelli et al., 2014; Huertas et al., 2000; Simionato et al., 2011). However, limitations such as low growth rate under stress conditions and high costs associated with downstream processing have hindered the commercialization of algae-based bio-fuel (Wijffels & Barbosa, 2010). Also, it is hard to emulate the abovementioned stress conditions in open raceway ponds for biofuel production.

As natural inhabitants of water eco-systems, which are prone to frequent and substantial variation in salinity, microalgae have adapted to accumulate particular metabolites and lipid under salinity stress. Significantly, salinity stress is easily achieved in open raceway ponds, and thus because of practical application in the field, we have pursued salinity over other stress conditions (Kim et al., 2016; Perrineau et al., 2014). Salinity-induced lipid accumulation has been investigated in several microalgal species, including halotolerant *Dunaliella* spp. (Takagi & Yoshida, 2006). Ho and colleagues (2017) gradually increased the salinity levels in the growth medium for *Chlamydomonas* sp. strain JSC4 and observed a lipid content of up to 59.4% of the biomass. Varying salinity levels induced lipid accumulation in *Nannochloropsis*, now *Microchloropsis* (see below) (Bartley et al., 2013; Gu et al., 2012). Under salinity stress, some marine euryhaline microalgae such as *Dunaliella tertiolecta* accumulate compatible solutes such as glycerol, which is of industrial importance (Goyal, 2007), indicating the potential for microalgae to be used for the production of high-value products in addition to lipid. Understanding the dynamics of gene expression during salinity stress may lead to the identification of high-value products that might be feasibly produced in addition to understanding how lipid production can be maximized. Recently, it was shown that strains with identical 18S rDNA sequences show considerable genomic variations (Corteggiani Carpinelli et al., 2014). Studies into the phylogeny of the genus *Nannochloropsis* using combined analysis of 18S rDNA and *rbc*L sequences indicated a new genus *Microchloropsis*, comprising the species *M. salina* and *M. gaditana*. *Microchloropsis gaditana* CCMP526, previously known as *Nannochloropsis gaditana* CCMP526 is one of the six algal species that was initially assigned to the genus *Nannochloropsis* (Fawley et al., 2015)*. Nannochloropsis* sp. is commonly found in brackish and ocean water. It is known for high eicosapentaenoic acid (EPA) and lipid content (Sukenik, 1991; Sukenik et al., 1989). Commonly used as a feed in the aquaculture industry, *M. gaditana* has been previously known to accumulate lipid (~50% dry cell weight) and carbohydrate in a variety of stress conditions.

The cellular response towards the acclimatization of *M. gaditana* towards hypersaline conditions was recently studied. Lipid accumulation increased by ~4.6 fold, while the carbohydrate content increased by ~1.7 fold (Karthikaichamy et al., 2018). Our earlier work has shown that the induction of lipid and carbohydrate accumulation occurred at different phases of growth in hypersaline conditions. The same phenomenon has also been observed in a strain of the freshwater alga *Chlamydomonas reinhardtii*, where starch was initially synthesized as a short-term carbon reserve, and then the cells switched to lipid accumulation for longer-term storage (Siaut et al., 2011). Lipid accumulation in hypersaline condition was earlier observed in *Acutodesmus obliquus*, *Chlorella vulgaris*, and in other *Nannochloropsis* species (Pandit et al., 2017). We systematically investigated the acclimatization in *M. gaditana* using RNA-Seq to observe genome-wide gene expression patterns in the genes involved in metabolic processes.

RNA-Seq relies on a suitably annotated reference genome sequence; in particular, genes involved in lipid accumulation should be annotated appropriately. RNA-Seq has been successfully used to reconstruct lipid and starch metabolic pathways in *Dunaliella tertiolecta* (Rismani-Yazdi et al., 2011), to identify the genes involved in terpenoid biosynthesis in *Botryococcus braunii* race B (Molnár et al., 2012) and to study triacylglycerol (TAG) accumulation in un-sequenced microalgae (*Neochloris oleoabundans* and *Chlorella vulgaris*) (Guarnieri et al., 2011; Rismani-Yazdi et al., 2012). An annotated draft genome sequence for *gaditana* strain B-31 is available, and this sequence has been used to map RNA-Seq read sets arising from the samples from the acclimatization study described in this chapter. The expression levels of annotated genes have been evaluated and compared under the various conditions and used to identify genes with significant differences in genes expression. The predicted functions attributed to these genes has enabled us to understand the metabolic pathways that have changed expression during acclimatization. Our findings support the previously observed physiological analysis and could pave the way for a better understanding of regulatory networks of major biosynthetic pathways for engineering stress resilient microalgal strains.

## Materials and methods

### Cultivation and harvesting

*Microchloropsis gaditana* CCMP526 was cultivated in 0.2 μm-filtered sea water (collected from the Gippsland Lakes, Gippsland, Victoria, Australia), supplemented with Guillard’s f/2 nutrients (Guillard, 1975) and 17 mM sodium nitrate, at 25°C using 500 ml glass bottles (Schott Duran, Germany). Cultures (300 ml) were mixed by bubbling with sterile air (0.2 μm filtered) supplied at a flow rate of 2.5 L min^−1^. Illumination was provided at 150 μmol · photons · m^−2^ · s^−1^ (Philips, TLD36W, Amsterdam, The Netherlands) with a light/dark cycle of 12/12 h. Sodium chloride was added to the existing f/2 media (control, 38 PSU) to make f/2 media of different salinities (55, 70 and 100 PSU). A portable refractometer (RHS-10ATC) was used to assess the salinity of the culture media. Cells at mid exponential phase were inoculated into f/2 media of different salinities at a concentration of approximately 4×10^6^ cells/ml. Cells were then sampled at specific time points (0, 1, 6, 24 and 72 h) that were chosen from our earlier physiological study (Karthikaichamy, 2018 #26). The samples were centrifuged (4000 xg, 4°C, 5 min) in an Heraeus Multifuge model 3SR Plus (thermos Scientific, Australia) and the resulting cell pellet was washed thrice with sterile distilled water to remove any residual salt from the growth medium. The pellet was then stored at −80°C until further processing.

### RNA isolation and quantification

RNA was isolated using Bioline II DNA/RNA/Protein extraction kit (Bioline, Australia) following the manufacturer’s protocol. The RNA samples were quantitated using the Invitrogen Qubit and associated chemistry, which incorporates a double-stranded DNA or RNA-specific fluorescent dye (Invitrogen, Carlsbad CA., USA).

### Quality control, library preparation and sequencing

The integrity of the samples was measured using an Agilent Bioanalyzer 2100 microfluidics device, in conjunction with the associated hardware and chemistry (Agilent Technologies, Waldbronn, Germany). Libraries were constructed for all the samples that passed the integrity test (RIN>8) using an Illumina TruSeq Stranded mRNA kit, following the manufacturer’s protocol. Roughly 800 ng of RNA was used for each library construction. The libraries were quantified using a Qubit DNA HS kit, which incorporates a double-stranded DNA-specific fluorescent dye (Invitrogen, Carlsbad CA., USA). Further, libraries were sized and checked for adapter contamination using the Agilent Bioanalyzer 2100 microfluidics device, in conjunction with Agilent DNA HS kits and chemistry (Agilent Technologies, Waldbronn, Germany). Libraries were then sequenced using Illumina NextSeq500 platform, producing single end, 75 base read-sets.

### Transcriptomic data analysis

The sequencing data were analysed with RNAsik pipeline (Tsyganov et al., 2018) using Ensembl reference files for *M. gaditana* strain B-31 (Accession number: GCA_000569095.1; FASTA and GTF) and differential gene expression analysis was performed using the Degust webtool (Powell, 2015), using ‘limma-voom’ for the statistical test. Statistical cut-off (p-value<0.01 and abs log_2_(fold change) >1) was applied to filter out the in-significant genes.

## Results and discussion

### Overview of transcriptomic data

The *M. gaditana* transcriptome analysis under high saline conditions was performed at two different salinity conditions (55 and 100 PSU) and at five different time points (0, 1, 6, 24 and 72 h) with sampling performed from triplicate independent cultures. The sampling time points, and salinity levels were identical to those used for the proteomics study. A total of 1170.5 million raw sequence reads was obtained from the RNA-Seq analysis for the 45 samples (average 26 million reads per sample). The reads were then mapped to the reference genome (Accession number: GCA_000569095.1). The genome of *M. gaditana* B-31 contains a higher number of predicted genes than strain CCMP526 and 94% of annotated CDS in CCMP526 are found in B-31 (Corteggiani Carpinelli et al., 2014). Therefore, the genome of *gaditana* B-31 was used as reference. Figure 1 shows the mapping statistics for all reads in each sample. On average ~71.4% of the reads were uniquely mapped and around 50.2% of the total reads were assigned to a gene, proportions normally seen at the RNA-Seq mapping stage (Ewels et al., 2016). The reads that were mapped to multiple loci were filtered out. The mean quality (Phred score) of the reads across all the samples was 34.6 which is well above the cut-off (27) (supplementary figure 1) and the adapter content in the reads were less than 0.32%, which suggests the absence of adapter dimers in the sequence library (supplementary figure 2). The salinity-induced transcriptome description of *M. gaditana* is listed in supplementary table S1.

**Figure 1.**
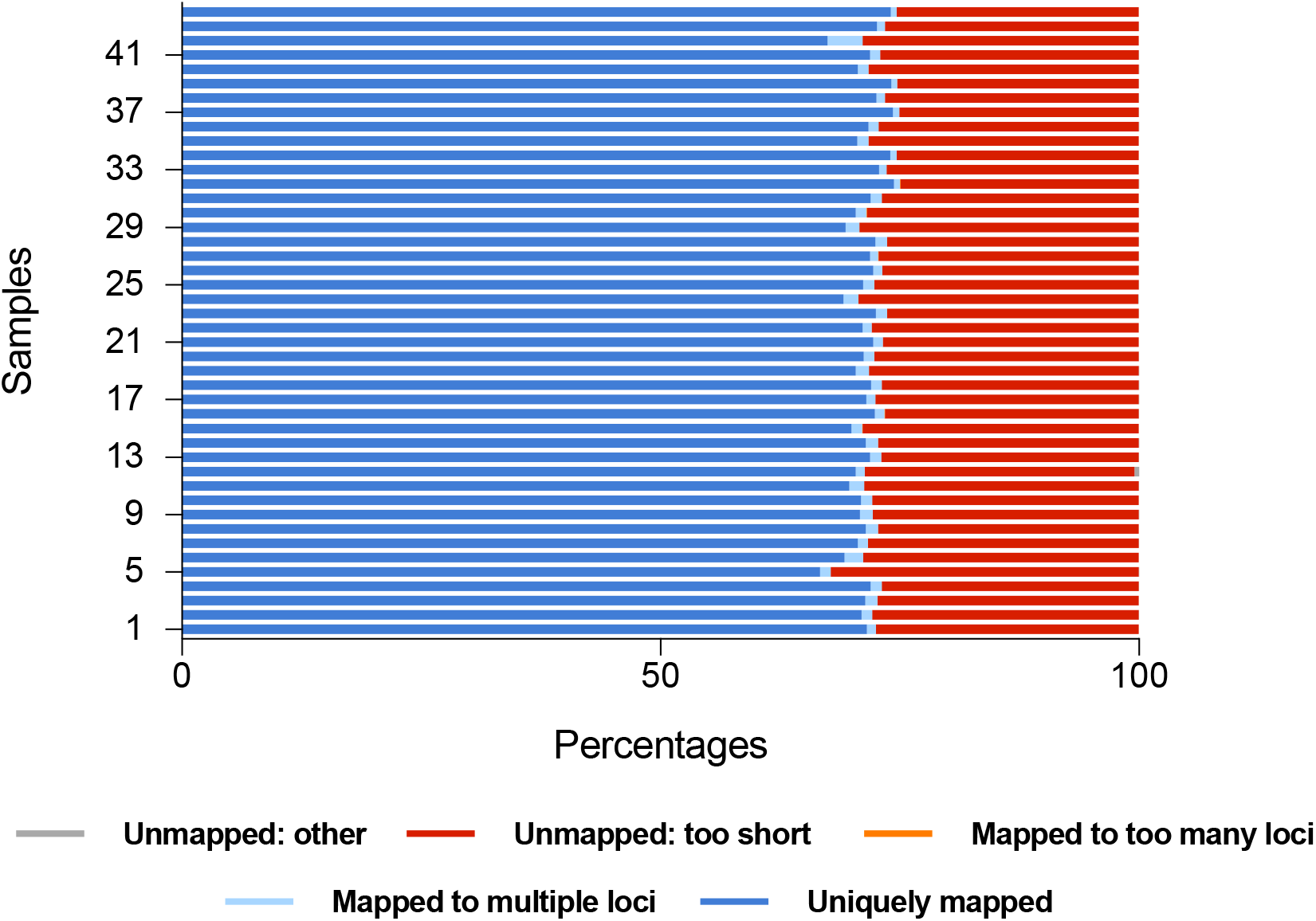
Mapping statistics of RNA-Seq reads (45 samples) of *M. gaditana* CCMP526 exposed to different levels of salinity at various time points.

To evaluate the differences and similarities of the transcriptomic data, the read-sets (triplicates) were reduced and plotted on a three-dimensional (3D) scale using multidimensional scaling (MDS) technique (figure 2). The greater the similarity between the samples, the closer the points representing the experimental conditions in the plot should be (Green et al., 1989). Clustering of data points indicate similarity between triplicate independent samples.

**Figure 2.**
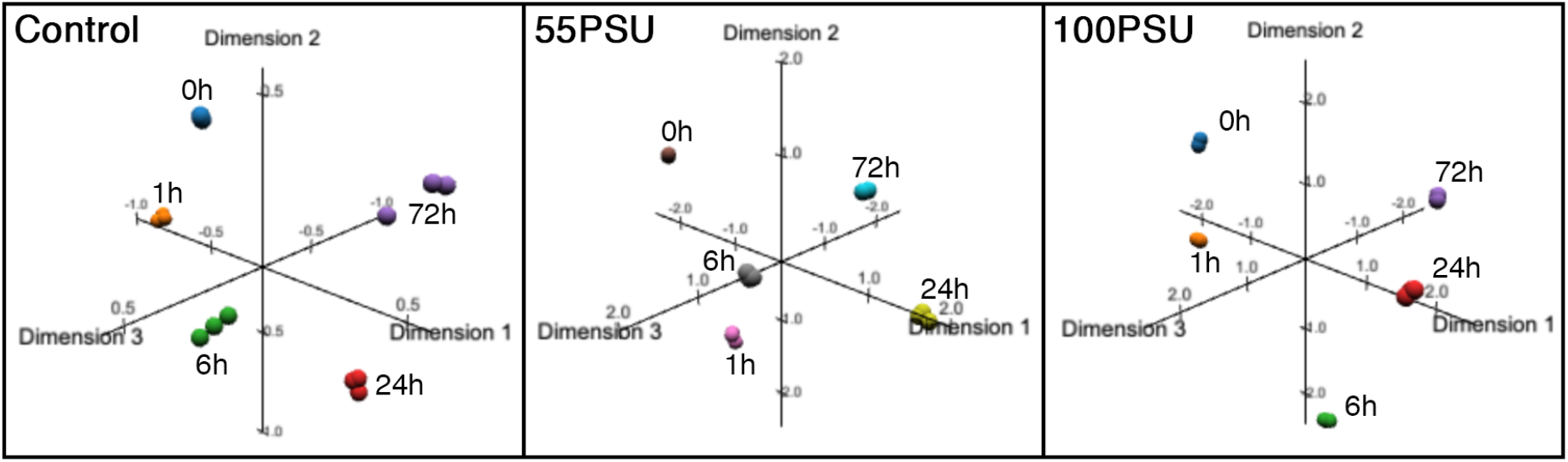
MDS plot of temporal RNA-Seq data of *M. gaditana* CCMP526 exposed to different levels of salinity.

The expression levels of the genes under control conditions (38 PSU) at each time point was normalised to 0 for the purpose of comparing expression levels of genes at 55 and 100 PSU. Filtering of differentially expressed genes were based on the following criteria,

- P-value should be less than 0.01 (p-value<0.01)
- Absolute log2(fold change) >1

From each datum point and salinity levels the genes that satisfied the stringent cut-off were selected for further analysis. Since we are studying the temporal gene expression patterns in hyper-saline conditions, genes that were significantly differentially expressed in at least one time point were selected across other time points too. We observed that at 55 PSU, the percentage of up-regulated genes was higher at 1 and 72 h and lower at other time points (0, 6 and 24 h), while in 100 PSU, the fraction of up-regulated genes was only higher at 1 h (figure 3).

**Figure 3.**
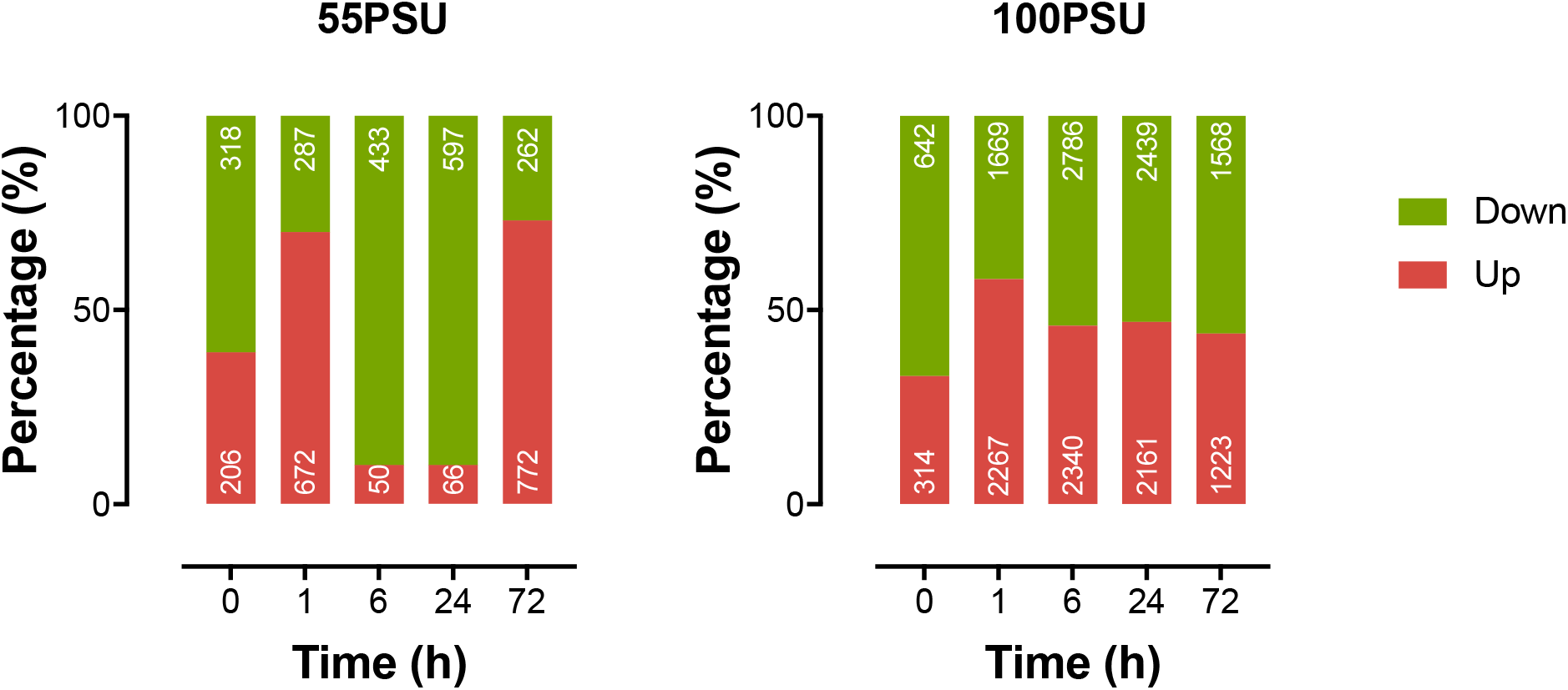
Percentage and number (in white) of differentially expressed genes in *Microchloropsis gaditana* CCMP526 under hyper-saline conditions.

We also observed that a higher number of genes were differentially expressed at 100 PSU than at 55 PSU. In total, 2522 and 6886 genes were differentially expressed at 55 PSU and 100 PSU respectively, out of which 2484 genes were differentially expressed at both conditions. Around 4402 genes were exclusively identified at 100 PSU, signifying the molecular state of adaptation of *M. gaditana* at high salt conditions. The genes exclusive to 100 PSU were predominantly involved in cell division, photosynthesis, translation initiation, lipid metabolism. It is also worthy to note that the genes for hypothetical proteins and uncharacterized proteins accounted for 41% and 37% in 55 and 100 PSU respectively. This could be due to the presence of novel biosynthetic pathway genes, that are not yet characterised, as mentioned in Corteggiani Carpinelli et al. (2014). A list of differentially expressed genes in *M. gaditana* in high saline conditions (55 and 100 PSU) is presented in supplementary tables S2 and S3.

### Salinity induced differentially expressed genes (DEGs)

Correspondence analysis (CA) is a multivariate method that is used to reduce the dimensions of a complex dataset into a low-dimensional space while preserving the information (Greenacre, 1984). CA has been widely used in exploring the interactions between the genes and experimental conditions (Fellenberg et al., 2001) or tissue types (Kishino & Waddell, 2000). By applying this method to our temporal transcriptomic data, we found that the first two axis accounts for 73.8% and 73.7% of the total inertia for 55 PSU and 100 PSU respectively. In the biplot (supplementary figure 3), each point represents a gene and the arrows that originate from origin represents time points. The genes that are closer or along the direction of the arrow are up-regulated at that particular time point and the genes that are on the opposite quadrant of the direction of the arrow represent the down-regulated genes at that particular time point. The genes are coloured according to their contribution towards the plot; highest contribution indicates that the gene has been highly differentially expressed along the time points.

The biplot for 55 PSU (supplementary figure 3) shows that the initial time points (0 and 1 h) and other time points (6, 24 and 72 h) are projected on the opposite side of each other of the first axis which accounts for maximum variation. While time points 0, 1 and 72 h are projected below zero, time points 6 and 24 h are projected above zero on second axis. This means that the first axis covers variation in the expression levels of the genes across the short and long-term responses. In the second axis, 72 h is projected along with the initial time points (0 and 1 h) signifying the acclimatisation path of *M. gaditana*, possibly through resuming the expression of genes projected below zero on second axis.

The short (0, 1 and 6 h) and long-term (24 and 72 h) associations are separated well along the first axis (42.1%). Most of the significantly expressed genes are projected below zero on the first axis, where 24 and 72 h are projected. Very few genes (15 out of 50) are associated with the initial responses (0, 1and 6 h). The second axis accounts for 31.6% of the total variation and it separates 6 and 72 h from 0, 1 and 24h time points. Interestingly, the projections of the time points for 55 and 100 PSU are not the same, implicating the different acclimatisation path of *M. gaditana* at different saline conditions.

Top 10 genes, based on their contribution values from each salinity conditions are plotted in figure 4. The top contributing genes in 55 PSU are mostly hypothetical protein coding genes, except Naga_1Chloroplast50 which codes for 50S ribosomal protein L6 (*rpl6*). This ribosomal protein binds directly to 23S ribosomal RNA and is located at the aminoacyl-tRNA binding site of the peptidyltransferase center. The GO terms are implicated in embryo development ending in seed dormancy and response to cytokinin in *A. thaliana* homologue (AT1G05190) (E value: 4e-53) (Černý et al., 2013). The expression of Naga_1Chloroplast50 is up-regulated by ~2-fold and ~8-fold at 6 and 72 h respectively indicating a role in alleviating salinity stress in *M. gaditana*. Interestingly, the hypothetical proteins were either localised in nucleus or chloroplast, signifying the role of this major organelle in response to hyper-saline conditions. An exception is Naga_100225g5, whose gene product is localised in mitochondria. A nucleus encoded hypothetical gene Naga_100040g27 is up-regulated during the initial phases of growth in hyper-saline condition (~32-fold and ~4-fold at 1 and 6 h respectively). The gene product of Naga_100040g27 is primarily composed of coils and has a PANTHER family of Late Embryogenesis Abundant Plants LEA-related (PTHR23241) (Mi et al., 2017). The Late Embryogenesis Abundant (McLean et al.) proteins are involved in response to drought and accumulate at the onset of seed desiccation in plant tissues (Olvera-Carrillo et al., 2011).

**Figure 4.**
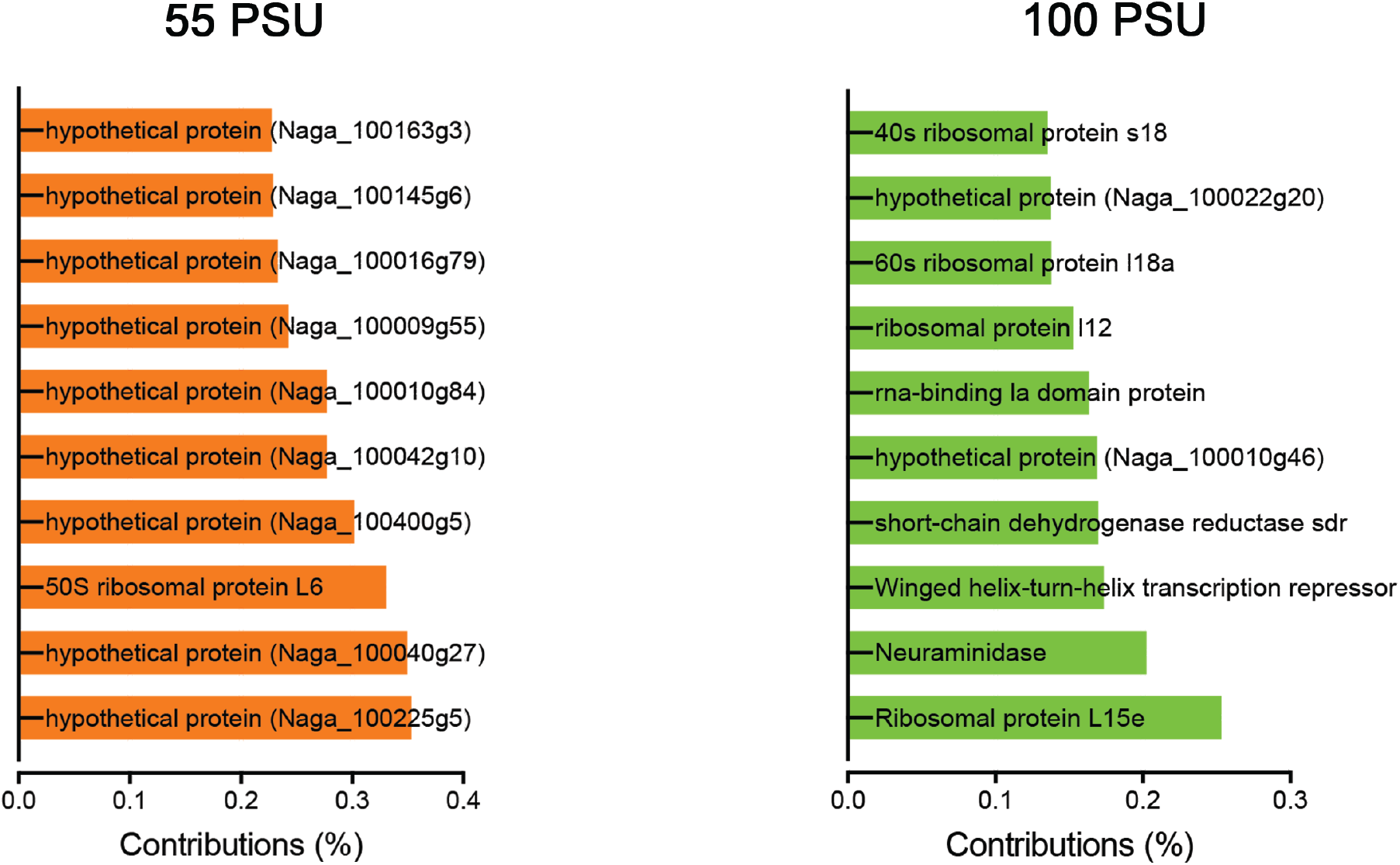
Correspondence analysis of differentially expressed genes in *M. gaditana* CCMP526 grown at 55 and 100 PSU. Top 10 gene contributors the contribute the dimensions in the biplot

The other hypothetical genes (Naga_100042g10 and Naga_100016g79) had homologues in *Tolypocladium ophioglossoides* CBS 100239 (a parasitic fungus) and *Lingula anatina* (marine organism, commonly known as ‘Lamp shell’). While the former is a Vacuolar amino acid transporter 3 gene that is involved in transport of amino acid efflux from vacuole to cytoplasm, the latter is spermatogenesis-associated protein 20 isoform X1, a thioredoxin domain-containing protein (IPR013766) which has active roles in redox regulation and defence against oxidative stress (Wei et al., 2012). The expression patterns of Naga_100042g10 and Naga_100016g79 are also similar (up-regulation during 0 and 1 h).

Contrarily, only two genes with products annotated as hypothetical proteins were found amongst the top contributors at 100 PSU. Other genes broadly fall into the category of genes that are involved in translation or a component of ribosome, implicating that the genes that code for the translation machinery is widely affected at 100 PSU. The ribosomal protein genes (Naga_100099g18, Naga_100003g20, Naga_100113g12) are highly up-regulated at 24h (up-to 64x) with an exception of Naga_100034g18 (60S ribosomal protein l18a), which was heavily down-regulated (~512x) through the growth curve. Similarly, Omidbakhshfard and colleagues have identified several translation-related genes in *Arabidopsis thaliana* to be affected by salinity (Omidbakhshfard et al., 2012). It could be proposed that synthesis of ribosomal proteins helps rebuilding the translational apparatus under salinity stress in *M. gaditana* as observed in maize (Zörb et al., 2004).

Neuraminidase is encoded by Naga_100045g1 in *M. gaditana* is significantly up-regulated at 24h (~512x) and down-regulated at all other time points (~6x and ~512x at 6 and 72 h respectively). Neuraminidase is predicted to be localised at cell wall and nucleus, characterised by WD40/YVTN repeat-like-containing domain superfamily (IPR015943) which is found in archaeal surface layer proteins (SLPs) that guard the cells from adverse environments (Jing et al., 2002). Another gene that is significantly differentially expressed at 100 PSU is Naga_100265g3 (short-chain dehydrogenase reductase, *sdr*), with the up-regulation (up-to ~1024x) at 6 and 72 h and down-regulation (~512x) at 24h. The short-chain dehydrogenase reductase is an oxidoreductase enzyme that belongs to a large family of enzymes, most of which are known to be NAD- or NADP- dependent oxidoreductases and possess a characteristic Rossmann fold (Rossmann et al., 1975). These enzymes are involved in wide range of functions ranging from stress response (Brosché & Strid, 1999) to secondary metabolite bio-synthesis (Sonawane et al., 2018).

### Clustering provides information on the temporal gene expression patterns

Expression patterns of significantly differentially expressed genes in each salinity levels (55 and 100 PSU) were grouped based on the optimum number of clusters estimated by figure of merit analysis (FOM) in Multiexperiment Viewer (Saeed et al., 2003). The FOM analysis was performed for 20 iterations and the optimum number of clusters was found to be seven for both 55 and 100 PSU (supplementary figure 4). GO-terms (biological process, cellular component and molecular function) were enriched (P-value<0.05) for each cluster and thus each cluster with a specific expression pattern is associated with the GO-terms. In 55 PSU, oxidoreductase activity was enriched in cluster2, where the 752 genes are up-regulated during the initial phases of growth (1 h). In cluster4 and 7, the genes are predominantly up-regulated during the later phases of growth (72 h). These clusters are enriched with cytoskeleton organisation, vesicle-mediated transport, translation and ribosome GO-terms. This implies that *M. gaditana* acclimatises to saline condition (55 PSU) by re-organising the cellular morphology and increasing the translation rate. The increase in biovolume of *M. gaditana* in 55 PSU at 72 h was observed in our previous physiological studies (Karthikaichamy et al., 2018). The short-term response includes GO-terms such as oxidoreductase activity and phosphatidylinositol binding (cluster 2 and 6) (figure 5). Earlier studies have shown that NaCl treatment induces the increase in the levels of phosphatidylinositol 4,5-bisphosphate and diacylglycerol pyrophosphate in *Arabidopsis thaliana* (Pical et al., 1999).

**Figure 5.**
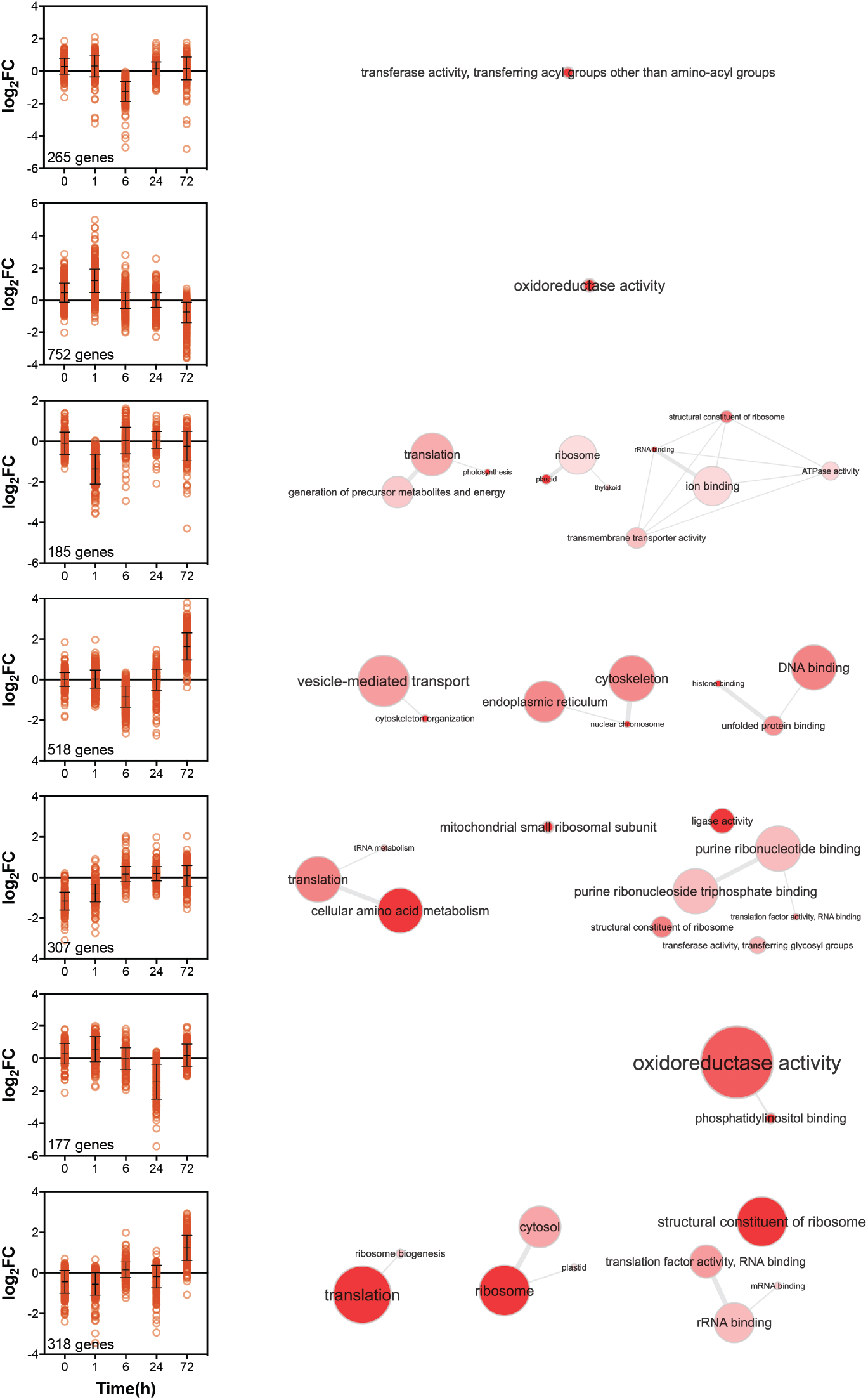
k-means clustering of significantly expressed genes in *M. gaditana* CCMP526 grown at 55 PSU. Interactive map of enriched GO-terms for corresponding cluster. Bubble color indicates the user-provided p-value; bubble size indicates the frequency of the GO term.

In 100 PSU, GO-terms such as cell division, photosynthesis, translation initiation, lipid metabolism, nucleus and thylakoid were enriched in cluster2, in which the genes are up-regulated at 24h. Though we observed a modest increase in lipid accumulation in 100 PSU only after 24h in our physiological studies (Karthikaichamy et al., 2018), the early up-regulation of genes involved in lipid metabolism can be attributed to the post-transcriptional dynamics, that is yet to be completely studied. Whereas, translation was down-regulated during the initial phases of growth (0, 1 and 6 h), as evident from cluster3. Comparable expression patterns of genes involved in translation was also shown by correspondence analysis, where the genes were down-regulated during the initial phases of growth in hyper-saline conditions (100 PSU). Cellular protein modification and fatty acid synthase complex were enriched in cluster1 during 1 and 6 h, but the p-value was greater than 0.05. Transmembrane transport and lyase activity were enriched at 6 and 72 h (figure 6), indicating the acclimatization strategy of *M. gaditana* at later phases of growth in hyper-saline conditions (100 PSU) by retaining the cytosolic K^+^/Na^+^ ratio or by maintaining cellular homeostasis by solute accumulation (Conde et al., 2011).

**Figure 6.**
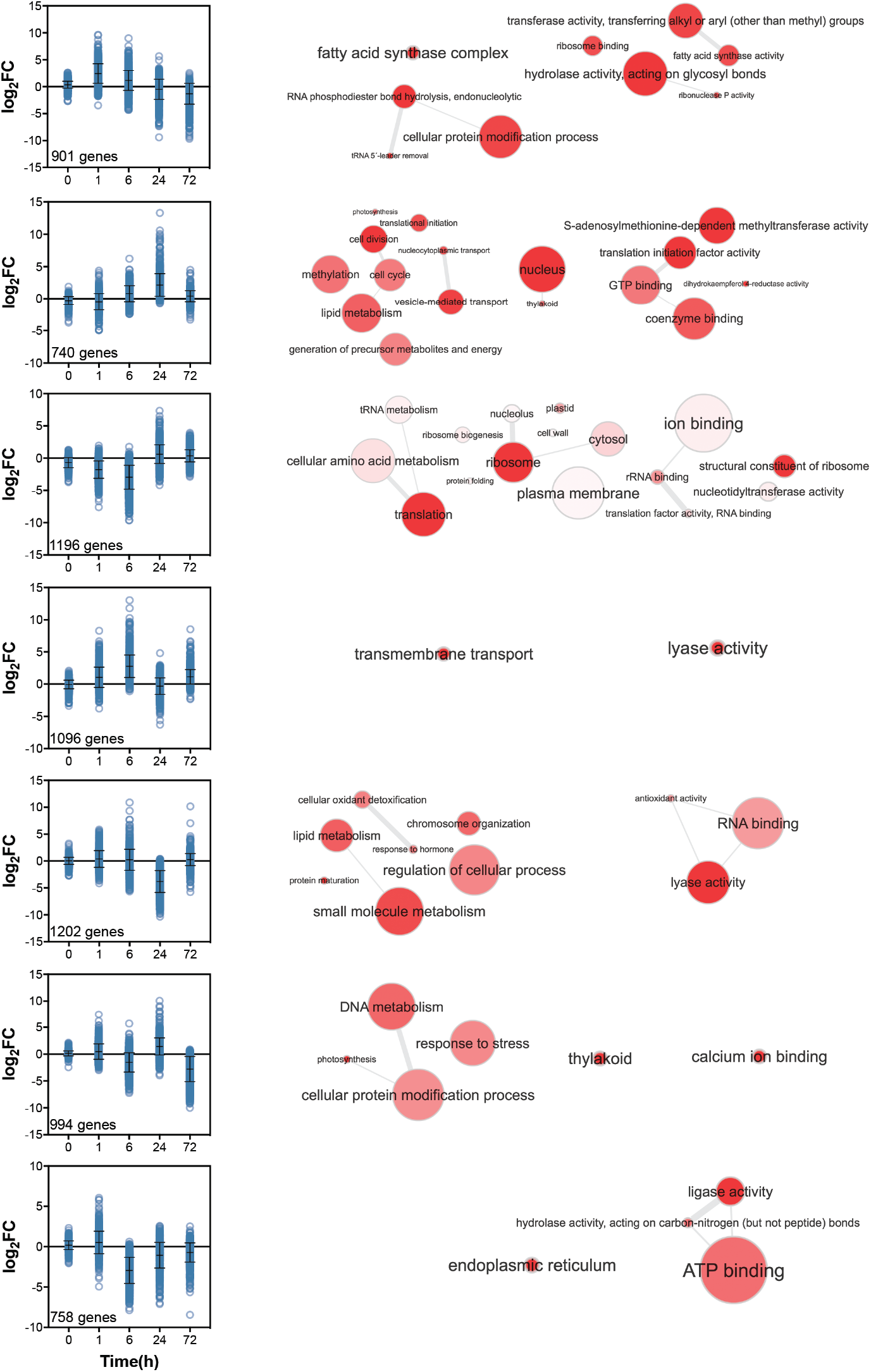
k-means clustering of significantly expressed genes in *M. gaditana* CCMP526 grown at 100 PSU. Interactive map of enriched GO-terms for corresponding cluster. Bubble color indicates the user-provided p-value; bubble size indicates the frequency of the GO term.

### Fate of cellular metabolism in high saline conditions

*Microchloropsis gaditana* CCMP526 employs a different strategy for acclimatization to different salinity levels that included cellular detoxification, lipid and carbohydrate biosynthesis (Karthikaichamy et al., 2018; Karthikaichamy et al., 2020). We also observed salinity induced transcriptome dynamics in *M. gaditana* emphasizing on lipid, carbohydrate metabolism and osmolyte biosynthesis. The significantly expressed genes from each time point were selected for pathway analysis and their biological significance during hyper-saline conditions are outlined in the following sections.

### Lipid metabolism is up-regulated in hyper-saline conditions

*Microchloropsis* accumulates high amounts of lipid during nutrient limited conditions (Bondioli et al., 2012). Lipid accumulation was observed in *Nannochloropsis* grown at different salinity levels (Bartley et al., 2013; Gu et al., 2012). Simionato and colleagues showed that TAG accumulation is primed by both *de novo* biosynthesis of fatty acids and degradation of membrane lipids in nitrate starved *Nannochloropsis gaditana* (Simionato et al., 2013). In our study, we have found several genes involved in lipid metabolism to be affected by high saline conditions (supplementary tables S4 and S5). In 55 PSU, several genes coding for lipid biosynthetic proteins acyl-synthetase (Naga_104758g1), diacylglycerol acyltransferase family protein (Naga_100006g86) and lipase, class 3 (Naga_100104g14) were found to be up-regulated at 1 h and 72 h by up-to 8x fold change (table S3.4). The gene Naga_100158g20, which encodes a long-chain acyl-synthetase (*LACS*) was up-regulated by 2-fold during the initial phases of growth (0 and 1 h). While in 100 PSU, an isoform of *LACS* (Naga_100014g59) was up-regulated by up-to 8-fold at 6 and 24h. *LACS* is essential for trafficking of lipid from endoplasmic reticulum (ER) to plastid (Jessen et al., 2015), while providing the starting material for very long-chain lipid biosynthesis in *Arabidopsis thaliana* (Jessen et al., 2015). Diacylglycerol acyltransferase (DGAT) catalyses the penultimate step of TAG biosynthetic pathway by transferring acyl moiety to diacyl-glycerol to form triacylglycerol (Courchesne et al., 2009) and is a crucial enzyme required for TAG biosynthesis in perturbed *Chlamydomonas* (Liu et al., 2016). Diacylglycerol acyltransferase type 2 (Naga_100343g3) was up-regulated 6.5-fold at 72 h, which correlate well with our physiological and proteomic analysis of salinity induced lipid accumulation in *M. gaditana* (Karthikaichamy et al., 2018; Karthikaichamy et al., 2020), where an increase in Nile red fluorescence and DGAT protein accumulation was observed.

Several class 3 lipases (Naga_100008g63, Naga_100171g1 and Naga_100104g14) and a fatty acid desaturase (Naga_100013g52) were up-regulated up-to ~16-fold at 1 and 6 h and up-to ~11-fold at 6 and 72 h respectively in 100 PSU. We also observed an early salinity responsive class 3 lipase (Naga_100043g24) in 100 PSU, that was up-regulated by ~56-fold at 1 h. Likewise, triglyceride lipase (Naga_100045g8), which specifically acts on TAGs is up-regulated by ~6.5-fold at 6 h. Lipase catalyses one of the two pathways for transfer of acyl chain from one lipid to another (Boyle et al., 2012) and is also involved in lipid degradation pathway in nitrogen starved *Chlamydomonas reinhartii* (Li et al., 2012b). An acylglycerol lipase (CrLIP1) from *Chlamydomonas reinhardtii* whose transcription is negatively correlated to lipid accumulation has been identified (Li et al., 2012a). Correspondingly, in our study, the transcriptional levels of class 3 lipases have also been found to be negatively correlated to lipid content (data not shown).

The primary source of NADPH in oleaginous micro-organisms for desaturation and elongation reactions of polyunsaturated fatty acid (PUFA) is malic enzyme (ME). Over-expression of ME in various photosynthetic micro-organisms has shown to increase neutral-lipid accumulation by 48.42% in *P. tricornutum* (Zhu et al., 2018), 2.5-fold increase in *M. circinelloides* (Zhang et al., 2007) and 3.2-fold increase in *Chlorella pyrenoidosa* (Xue et al., 2016). However, in our study, there was a significant down-regulation of malic enzyme (Naga_100957g1) transcripts by ~8 to 12-folds. Physiological analysis of *M. gaditana* at hyper-saline conditions reveal that the neutral-lipid accumulates after 24h in 100 PSU (Karthikaichamy et al., 2018). Therefore, the cell depends on other sources of reducing equivalents, which is likely supplied by the pentose phosphate pathway. A similar condition in which the reducing equivalents (NADPH) were supplied via pentose phosphate pathway was observed in oleaginous yeast *Yarrowia lipolytica* (Wasylenko et al., 2015). Fold changes for representative genes are shown in figure 7.

**Figure 7.**
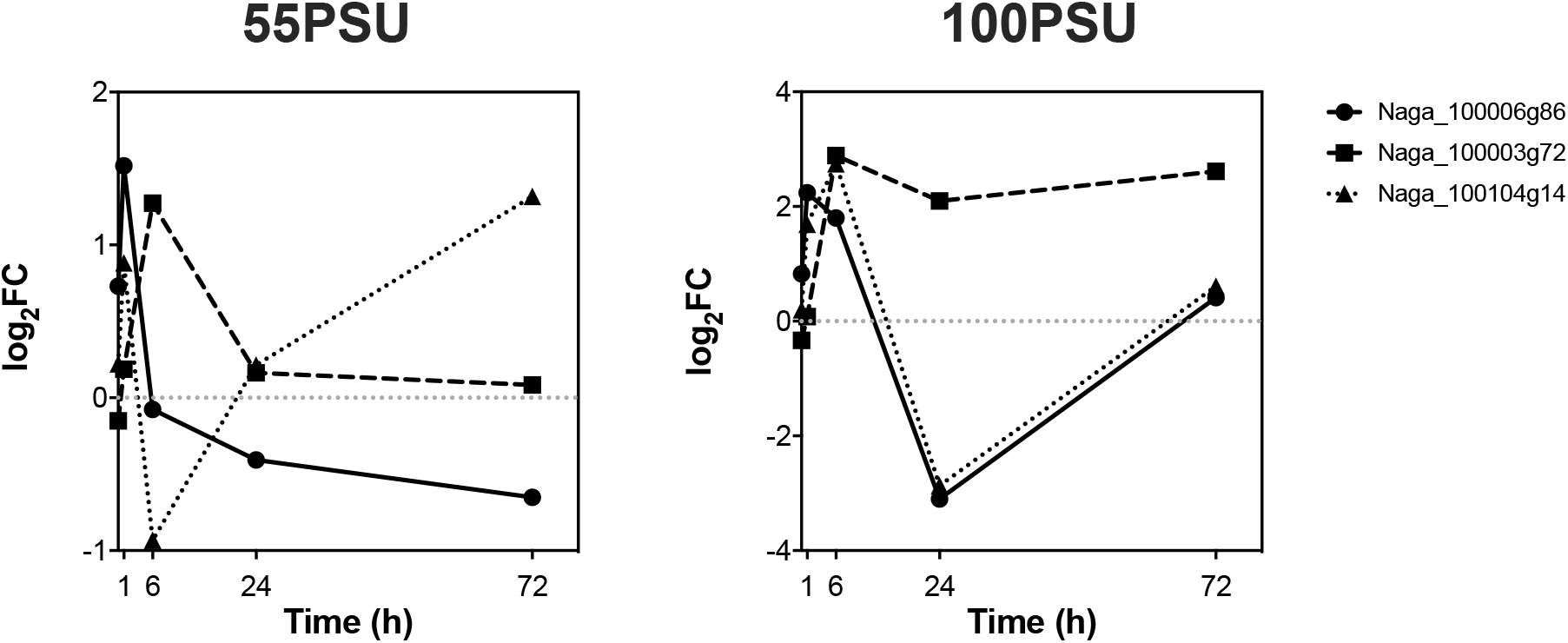
Temporal transcriptional changes of representative genes in lipid (Naga_100006g86 - Diacylglycerol acyltransferase family protein and Naga_100104g14 - lipase, class 3) and sterol (Naga_100003g72 - 24-dehydrocholesterol reductase) metabolism in *M. gaditana* CCMP526 in high-saline conditions (55 and 100 PSU).

### Membrane re-modelling mitigates salinity stress

The fluid nature of the plant and algal cell membrane can be altered by changing the lipid composition of the membrane by fatty acid desaturase. This phenomenon is beneficial for the organism to adapt to the stress conditions (Wada et al., 1994). The unsaturated fatty acids has also been indicated as the precursors of signalling molecules in plant defence mechanism (Kachroo & Kachroo, 2009). While the biovolume of *M. gaditana* at 72 h in 100 PSU significantly increased than the preceding time point (48h), fatty acid desaturase type 2 (Naga_100013g52) was also up-regulated signifying the importance of desaturation of membrane lipid for maintaining the cellular morphology during high saline conditions. Sterols are a part of lipid compounds that is used to regulate permeability and fluidity of membrane (Silvestro et al., 2013). Another desaturating enzyme 24-dehydrocholesterol reductase (Naga_100003g72) which catalyses the conversion of desmosterol to cholesterol is also found to be up-regulated ~8-fold at 6, 24 and 72 h in 100 PSU. While a modest up-regulation of ~2.5-fold was observed at 6 h in 55 PSU. The enzyme 24-dehydrocholesterol reductase (Naga_100003g72) is involved in the final steps of post-squalene sterol pathway to yield Δ^5^-sterols in wild-type *Arabidopsis thaliana* (Schaller, 2010). Sterols act as structural components of membrane that confer fluidity and permeability (Demel & De Kruyff, 1976; Yeagle, 1991). They also protect the cell during adverse conditions such as drought and heat shock (Beck et al., 2007; Posé et al., 2009). It is thus hypothesised that the combined effect of fatty acid desaturase and 24-dehydrocholesterol reductase benefits *M. gaditana* in overcoming the membrane stress in hyper-saline conditions by presumably increasing the membrane fluidity and acting as signalling molecules.

### Transcriptional regulation of carbon assimilation in *M. gaditana* under hyper-saline conditions

Ribulose-1,5-bisphosphate carboxylase/oxygenase (Rubisco) catalyses the rate-limiting step in CO_2_ fixation during photosynthesis (Spreitzer, 2003). The two subunits of Rubisco*, rbcS* (Naga_1Chloroplast92) and *rbcL* (Naga_1Chloroplast16) were transcriptionally down-regulated (~4x) at 1 h and then showed a non-significant recovery during the later phases of growth in 55 PSU (<1-fold change). In contrast, in 100 PSU, the CO_2_ fixing genes were transcriptionally down-regulated along the growth curve. Carbon dioxide fixation in eukaryotic photosynthetic organisms takes place in the chloroplast, in which the carbon is incorporated into organic molecules such as sugar phosphates, starch and a range of other compounds (Takabe et al., 1988). However, carbonic anhydrase (Naga_100024g5) which catalyses the reversible hydration of CO_2_ (Coleman, 2000) was up-regulated (~4x) during the initial (1 and 6 h) and later phase (72 h) of growth in 100 PSU. In photosynthetic micro-organisms and higher plants carbonic anhydrase converts HCO_3_^−^ to CO_2_ which is then fixed by ribulose 1,5-bisphosphate carboxylase/oxygenase (Espie & Kimber, 2011). Transcriptional up-regulation of carbonic anhydrase was observed in maize (Kravchik & Bernstein, 2013). Similarly, Yu and colleagues observed an increase in chloroplastic carbonic anhydrase mRNA in salt-treated *Arabidopsis* (Yu et al., 2007).

The activity and abundance of the two enzymes involved in carbon acquisition (Rubisco and carbonic anhydrase) are regulated by several abiotic factors such as nitrogen availability, CO_2_ concentration, temperature and light (Cornwall et al., 2015; Helbling et al., 2011; Ihnken et al., 2010; Madsen & Baattrup-Pedersen, 1995; Reed & Graham, 1981). Up-regulation of external carbonic anhydrase activity is induced by hyper-saline conditions in halotolerant *D. salina* as dissolved CO_2_ levels decrease with increasing salinity (Booth and Beardall, 1991). There exists a fine balance between Rubisco and carbonic anhydrase in allocation of CO_2_ and optimising photosynthesis (Tortell et al., 2006). Recently, a positive correlation was observed between Rubisco and carbonic anhydrase activities in *P. tricornutum*, indicating mutual dependency of the two enzymes in CO_2_ fixation (Zeng et al., 2019). Contrarily, we observed a negative correlation between the transcript abundances of Rubisco and carbonic anhydrase. While the mRNA levels of carbonic anhydrase (Naga_100024g5) were up-regulated (~4x), Rubisco transcripts (Naga_1Chloroplast92 and Naga_1Chloroplast16) were down-regulated (~4x) during the initial and later phases of growth in high saline conditions (100 PSU). The increase in carbonic anhydrase expression (protein product predicted to be localized in chloroplast and nucleus) is consistent with a possible up-regulation of CO_2_ concentrating mechanisms, as would be required by the lower concentration of CO_2_ in solution at elevated salinity (Booth and Beardall, 1991). The decreased expression of Rubisco genes is also consistent with the large decrease in photosynthetic capacity (as determined by maximum rates of electron transport) reported in *M. gaditana* at high salinity (Karthikaichamy, 2018 #46). Our findings suggests that *M. gaditana* invests in carbon acquisition than *de novo* Rubisco biosynthesis, as Rubisco biosynthesis requires a high N cost owing to its large size (550 kDa) and engagement of additional proteins for assembly and activation (Andersson, 2008 #153). Relative fold changes to control (38PSU) for representative gene transcripts involved in carbon acquisition are shown in figure 8.

**Figure 8.**
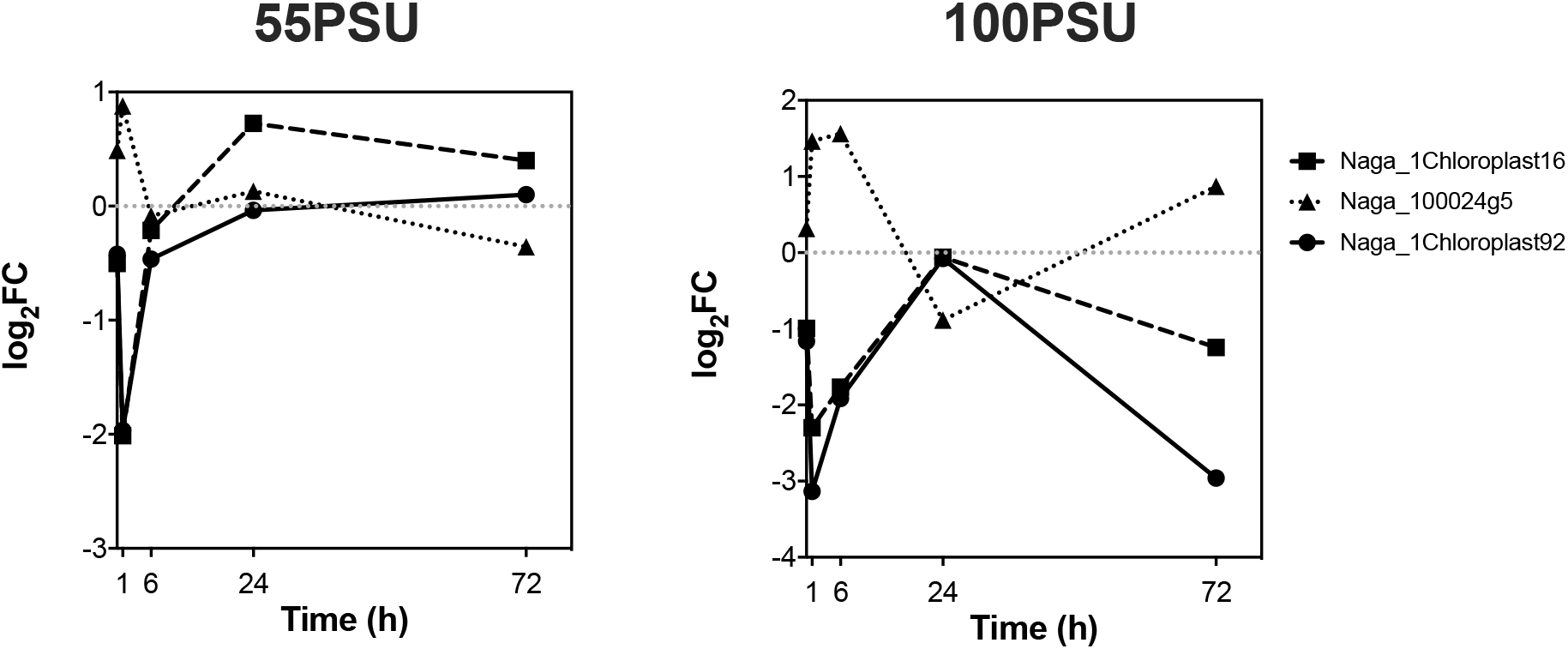
Temporal transcriptional changes of *rbcS* (Naga_1Chloroplast92), *rbcL* (Naga_1Chloroplast16) and carbonic anhydrase (Naga_100024g5) in response to increase in salinity levels in *M. gaditana* CCMP526. Relative transcript levels of carbonic anhydrase (Naga_100024g5) were insignificant in 55 PSU.

### Up-regulation of osmolyte biosynthesis genes in *M. gaditana* under hyper-saline conditions

One of the principal adaptive mechanisms in salinity-challenged halophytes is the production of osmolytes (Flowers & Colmer, 2008). Primarily accumulated in the cytoplasm, the osmolytes are highly soluble low-molecular-weight organic compounds that can range from amino acids, tertiary sulphonium compounds, quaternary ammonium compounds, sugars to polyhydric alcohols (Flowers et al., 1977; Wyn Jones, 1977). They protect cellular organelles and alleviate oxidative damage caused by abiotic stresses. Some osmolytes also confer stress tolerance by scavenging free radicals and protecting key enzymes (Szabados et al., 2011). It has been previously reported that about 20% of assimilated carbon is partitioned into glycerol in salinity-challenged *Dunaliella*, a halotolerant green alga (Craigie & McLachlan, 1964; Wegmann, 1971).

Consistent up-regulation in transcript abundance (up to 40x) of glycerol-3-phosphate dehydrogenase 1-like protein (GPD1) (Naga_100042g20) in 100 PSU was observed at all time points (1, 6, 24, 72 h) except at 0 h. Likewise, a modest up-regulation of Naga_100042g20 transcript was observed at 0 and 1 h (up to 8x) in 55 PSU. GPD1 is located in the mitochondrial inner-membrane space or cytosol and catalyses the reduction of dihydroxyacetone phosphate into glycerol-3-phosphate. GPD1 links carbohydrate and lipid metabolism by providing the glycerol backbone for TAG (Ou et al., 2006). Additionally, other genes involved in biosynthesis of compatible solutes for osmo-protection were also up-regulated at high saline conditions. For instance, choline monooxygenase (Naga_100113g3) and betaine aldehyde dehydrogenase (Naga_100034g16) are genes involved in the biosynthesis of glycine betaine (betaine) from choline. Choline monooxygenase and betaine aldehyde dehydrogenase catalyse the first and final step of betaine biosynthesis respectively. *Arabidopsis halophytica* cells respond to variety of salinity levels by fixing photosynthetic CO_2_ into betaine (Takabe et al., 1988). Exogenous addition of betaine to the growth medium of *A. thaliana* conferred tolerance to abiotic stresses (Hibino et al., 2002). Glycine betaine is known to accumulate in marine phytoplankton and macroalgae. It has also been implicated in accelerated recovery of PSII in *Synechococcus* (Deshnium, 1997 #154) and protection of cyanobacterial PSII complex during high saline conditions (Papageorgiou, 1995 #155).

Proline is another osmolyte used by plants and algae to combat stress, and studies have shown a positive correlation between proline accumulation and stress induction in plants (Kemble & Macpherson, 1954). Proline accumulation has also been implicated in responses to stress induced by drought (Choudhary et al., 2005), UV radiation (Saradhi et al., 1995), hyper salinity (Yoshiba et al., 1995) and heavy metals (Schat et al., 1997). Additionally, in microalgae e.g., *Anacystis nidulans*, *Chlorella sp*. and *Chlorella vulgaris*, proline accumulation was observed in exposure to Cu (copper) stress (Mehta, 1999 #158;Wu, 1995 #156;Wu, 1998 #157). In higher plants, proline biosynthesis uses glutamate as the precursor metabolite, which is then reduced to glutamate-semialdehyde (GSA) by the enzyme pyrroline-5-carboxylate synthetase (P5CS). GSA is spontaneously converted to a pyrroline-5-carboxylate (P5C) intermediate (Hu et al., 1992). Pyrroline-5-carboxylate reductase (P5CR) catalyses the final reduction of the P5C intermediate to proline (Verbruggen et al., 1993). In *M. gaditana*, one of the P5CR isoforms is encoded by Naga_100026g57 and showed significant increase in transcriptional abundance (~8x) during the initial phases of growth (0, 1 h) and down-regulation of transcripts at 6 and 24h (~10x) in 100 PSU, whereas *M. gaditana* grown in 55 PSU only showed a modest increase in Naga_100026g57 transcript at 1 h (~3x) post salinity stress. A significant increase in DCF fluorescence, indicative of the presence of reactive oxygen species, was observed at 24h in 100 PSU (Karthikaichamy et al., 2018), which is in accordance with the reduced transcript levels of P5CR (Naga_100026g57). Comparable results for P5CR overexpression were also observed in microalgae (*C. reinhardtii*) and plants where the free radical levels were reduced due to hyperaccumulation of proline (Siripornadulsil et al., 2002; Yamada et al., 2005). Relative fold changes to control (38PSU) for representative gene transcripts involved in osmolyte biosynthesis are shown in figure 9.

**Figure 9.**
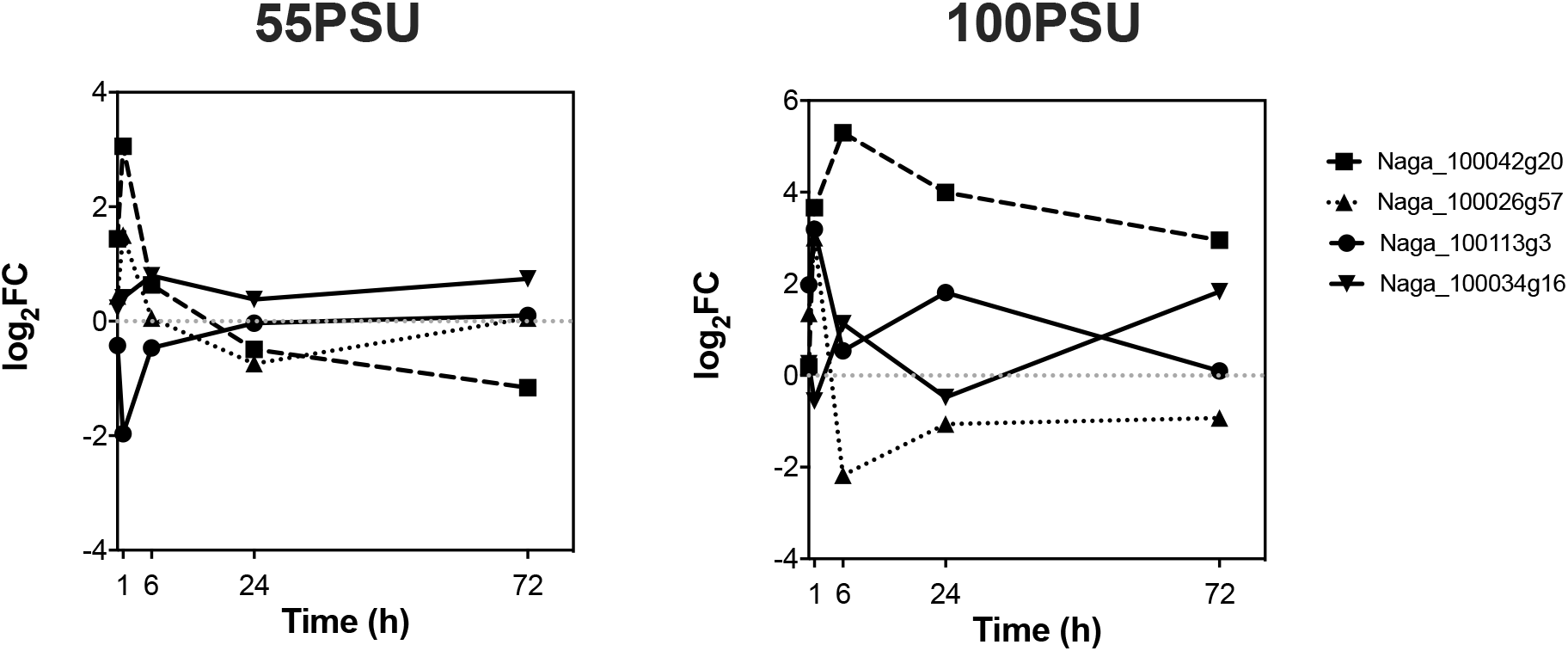
Temporal transcriptional changes in expression of genes for choline monooxygenase (Naga_100113g3), pyrroline-5-carboxylate reductase (Naga_100026g57), betaine aldehyde dehydrogenase (Naga_100034g16) and glycerol-3-phosphate dehydrogenase 1-like protein (GPD1) (Naga_100042g20) in response to increase in salinity levels in *M. gaditana* CCMP526. Relative transcript levels of betaine aldehyde dehydrogenase (Naga_100034g16) were insignificant in 55 PSU.

### Genes involved in redirection of carbon flux towards lipid biosynthesis

Interconversion of lipid and carbohydrates is inevitable during several stress conditions as the cell tries to shunt the carbon reserves towards biosynthesis of different biomolecules (Johnson & Alric, 2013). For instance, lipid accumulation was tightly linked with a decrease in pyrenoids (mostly starch and proteins) under phosphate-limited condition in *Scenedesmus* sp. (Yang et al., 2018). In *M. gaditana*, we observed a negative correlation between neutral-lipid accumulation and carbohydrate content in hyper-saline conditions (100 PSU) (Karthikaichamy et al., 2018). Therefore, the expression levels of the genes involved in carbohydrate biosynthetic pathway were assessed under exposure to hyper-saline conditions (55 and 100 PSU). Glucan endo-β-glucosidase (Naga_100464g2) catalyses the hydrolysis of complex polysaccharides (glucoside, cellulose, etc.,) to simple sugars such as glucose. The increased abundance of glucan endo-β-glucosidase results in a reduction of the carbohydrate content in the cell. The hydrolysed carbohydrates (glucose) are then converted into acetyl-CoA, a precursor for fatty acid synthesis. Abundance of the Naga_100464g2 transcript was significantly down-regulated (~3x) during the initial phases of growth (0 and 1 h) in 55 PSU, whereas, higher saline conditions (100 PSU) induced a down-regulation of this transcript. There was a modest down-regulation (~8x) of Naga_100464g2 during the initial phases of growth (0 and 1 h), and a strong down-regulation (~64x) was observed at 6 and 24h in 100 PSU.

Likewise, a similar trend was observed with the carbohydrate and neutral-lipid content in *M. gaditana* at high saline conditions. Carbohydrate content was at maximum (at 48 and 72 h) after the strong down-regulation of Naga_100464g2 at 6 and 24h in 100 PSU. Conversely, neutral-lipid content increased as the carbohydrate content decreased (at 72 h) in 100 PSU, where there was no significant change in the transcript levels of Naga_100464g2. This suggests that the up-regulation of the genes involved in carbohydrate catabolism could enhance lipid accumulation. Meanwhile, a gene encoding cellulose synthase (Naga_100079g22) was down-regulated in 55 PSU (around 8x at 24h) and 100 PSU (up to 64x; except 0 h). This suggests that carbohydrate synthesis was supressed at high saline conditions (55 and 100 PSU). The gene encoding the precursor biosynthesis of cellulose, UDP-glucose was also down-regulated at 100 PSU in *M. gaditana* (Naga_100013g89). UDP-glucose pyrophosphorylase 2 catalyses the synthesis of UDP-glucose from glucose, and down-regulation causes accumulation of glucose, which can be converted into acetyl-CoA for lipid biosynthesis.

Additionally, a gene encoding D-lactate dehydrogenase (Naga_100083g18) which catalyses the interconversion of lactate to pyruvate was significantly up-regulated throughout the growth phase (except 0 h) in 100 PSU though only at 1 h in 55 PSU. Maximum up-regulation of Naga_100083g18 occurred under 100 PSU, 24 h post transfer into the saline condition (~16x) and then the transcripts were modestly up-regulated by ~3x at 72 h. These findings are consistent with the decrease in carbohydrate content and an increase in lipid content after 24 h in 100 PSU (Karthikaichamy et al., 2018). Transgenic *Oryza sativa* with a silenced lactate dehydrogenase gene (*Osd-LDH* RNAi) was sensitive to salinity stress (An et al., 2017). Comparably, pyruvate kinase (Naga_100072g23) was also found to be significantly up-regulated during the later phases (24 and 72 h) of growth in high saline conditions (100 PSU). Though there was a significant down-regulation of Naga_100072g23 during the initial phases (0, 1 and 6 h) of growth in 100 PSU, we observed an up-regulation of ~8x at 24 and 72 h.

Relative fold changes to control (38PSU) for representative gene transcripts involved in carbohydrate biosynthesis are shown in figure 10. Choreographed up-regulation of D-lactate dehydrogenase and pyruvate kinase at high-saline conditions could result in accumulation of pyruvate, which could be shunted towards lipid biosynthesis via acetyl-CoA. A similar phenomenon was observed in phosphorus-starved *Scenedesmus sp.* (Yang et al., 2018).

**Figure 10.**
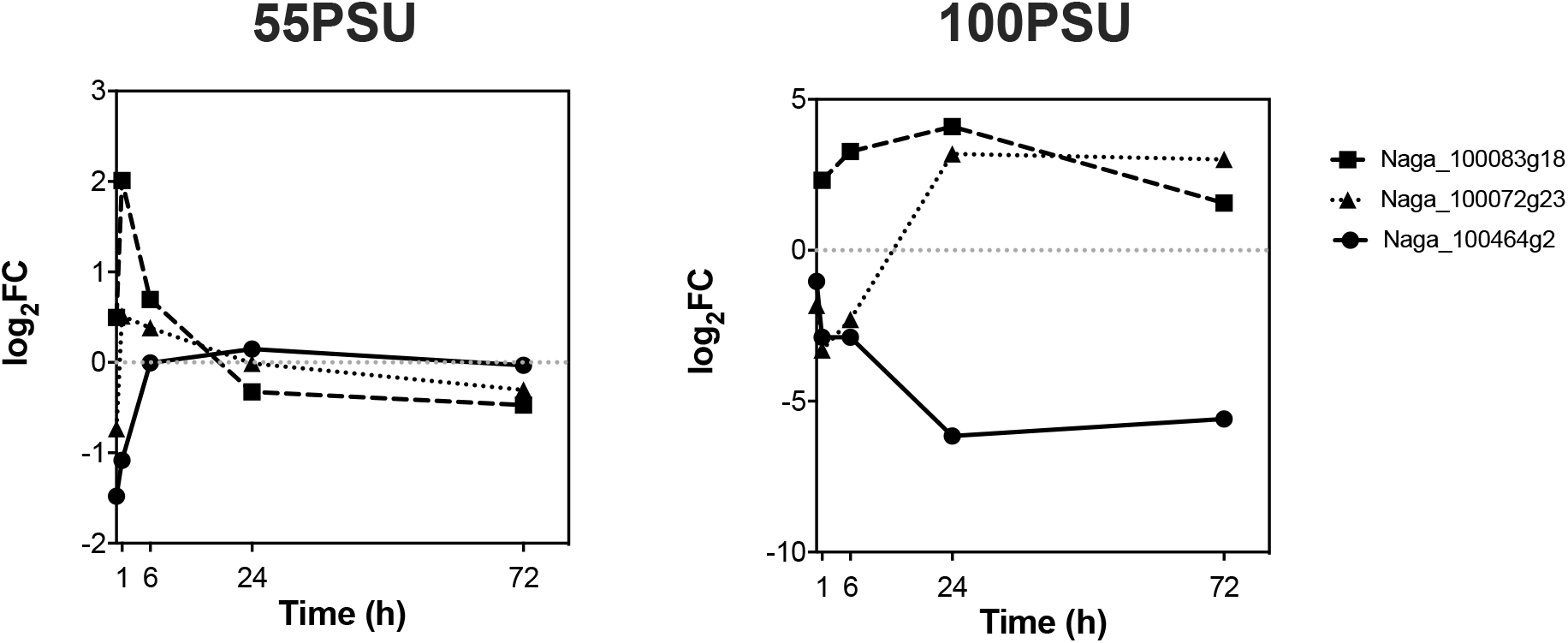
Temporal transcriptional changes of glucan endo-β-glucosidase (Naga_100464g2), D-lactate dehydrogenase (Naga_100083g18) and pyruvate kinase (Naga_100072g23) in response to increase in salinity levels in *M. gaditana* CCMP526.

### Conclusions

RNA-Seq was used to study the temporal transcriptomic response of *M. gaditana* to high salt conditions (55 and 100 PSU). Significant differentially expressed genes were clustered based on the expression pattern and GO-enrichment was performed to understand the dynamics of different bio-synthetic pathways under salinity stress. The major genes that responded to varying levels of salinity and growth period were investigated using correspondence analysis. Exposure to 100 PSU had a profound effect on the number of differentially expressed genes. This could be attributed to alterations in cellular morphology and metabolism. Genes involved in lipid metabolism, osmolyte biosynthesis, carbon acquisition and carbohydrate catabolism pathways were up-regulated in high saline condition. Our findings focus on several biosynthetic process that ultimately aid in acclimatization of *M. gaditana* in high saline conditions. The acclimatization process involves interplay of genes involved in carbon backbone assimilation and shunting to various biomolecules (lipid, carbohydrate and osmolytes). Also, the GO analysis of temporal RNA-Seq indicates *M. gaditana* employs a varied response to different salinity levels. Finally, transcriptome profiling of oleaginous microalga *M. gaditana* under hyper-saline conditions was used to understand the dynamics and inter-play of genes involved in various biosynthetic pathways. In future, gene expression profile can be used to identify up-stream regulatory elements for synthetic biology applications and also to develop salinity tolerant microalgae.

## Supplementary information

### Supplementary figures

**Supplementary figure 1.**
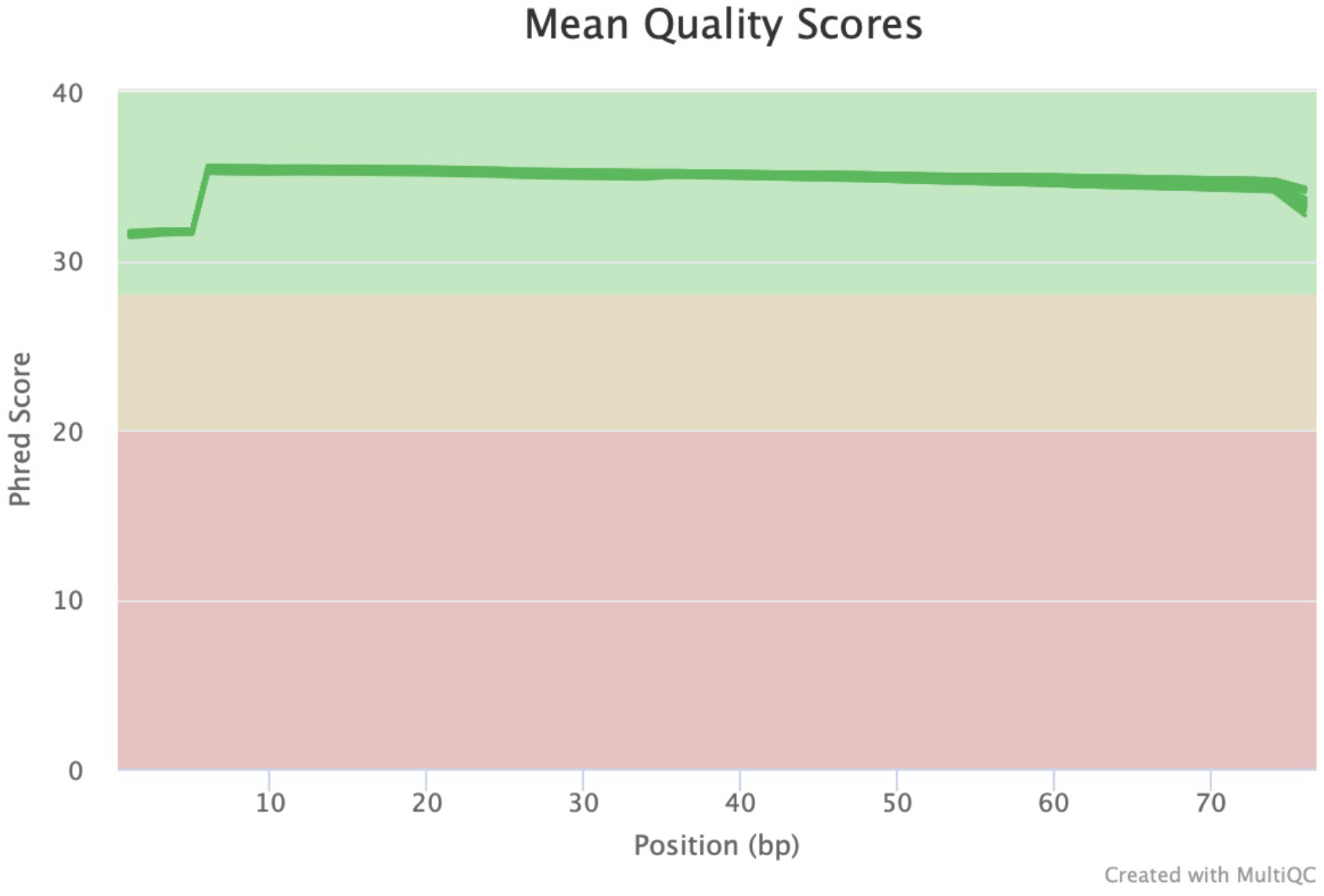
Phred quality score of the average distribution of reads across all samples.

**Supplementary figure 2.**
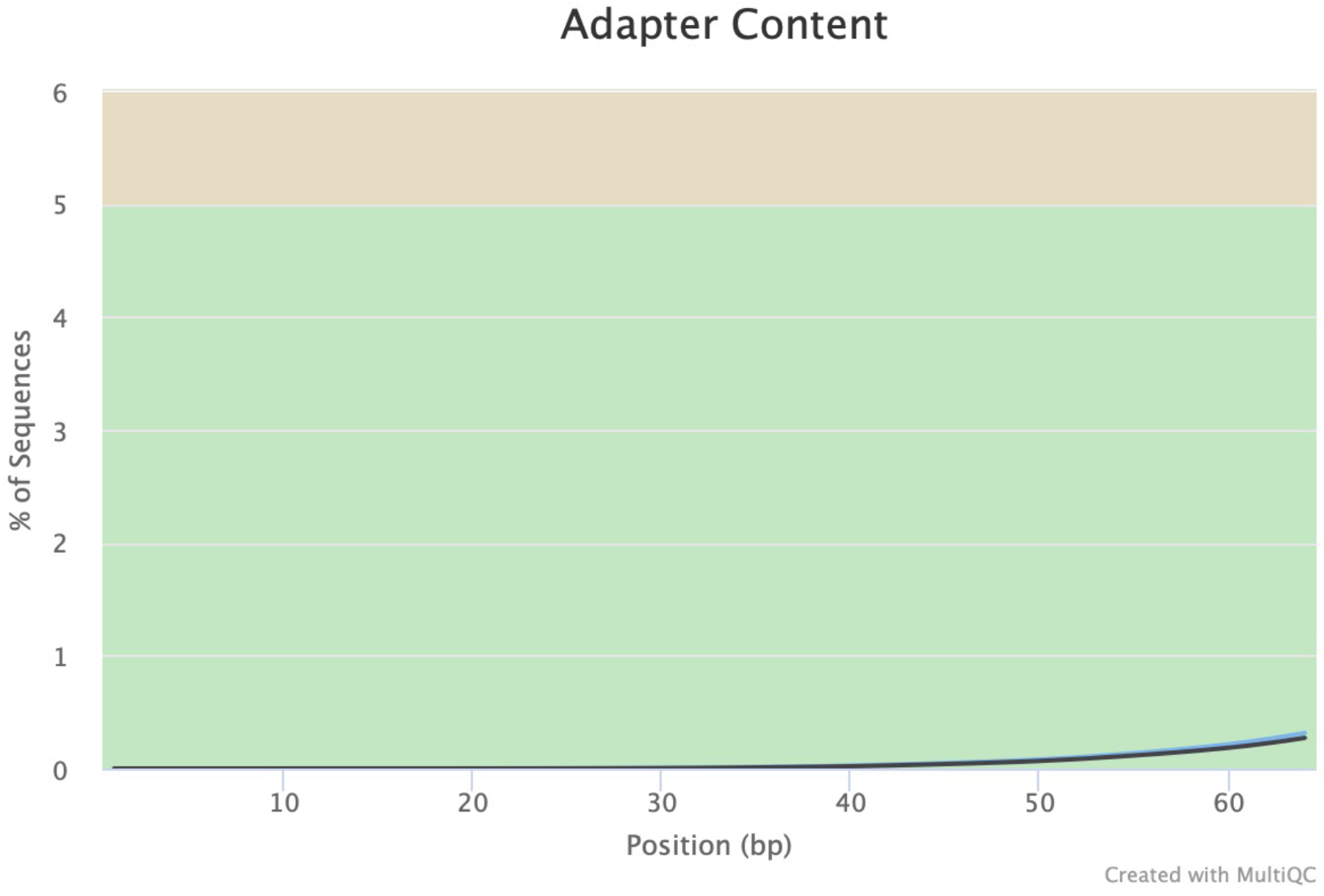
The cumulative percentage count of the proportion of the adapter sequences at each position in RNA-Seq library (samples with ≥ 0.1% adapter contamination are displayed).

**Supplementary figure 3.**
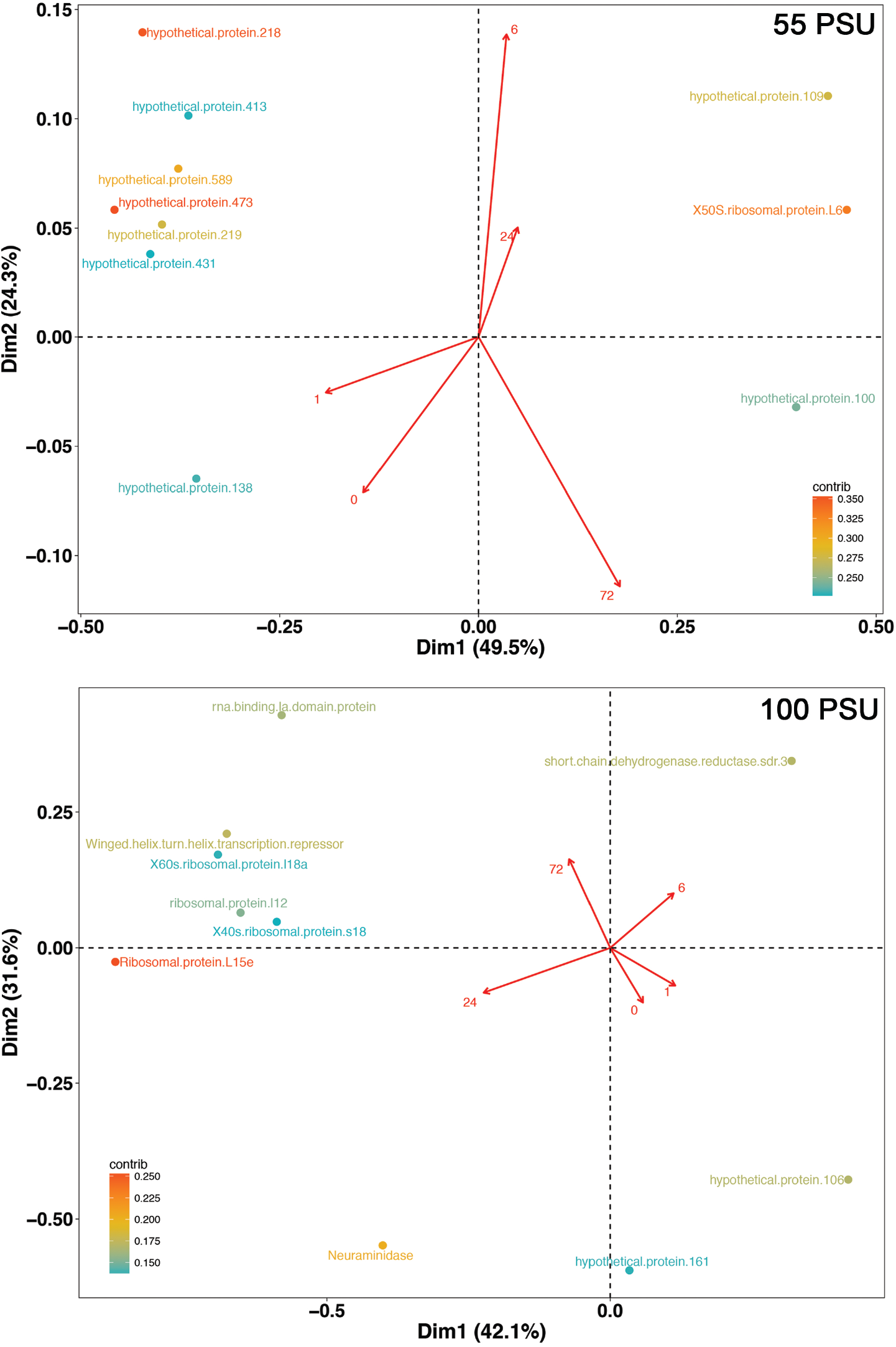
Correspondence analysis of differentially expressed genes in *M. gaditana* CCMP526 grown at 55 and 100 PSU. Biplot showing the association between time points and differentially expressed genes (numbers along the red arrow represent time points).

**Supplementary figure 4.**
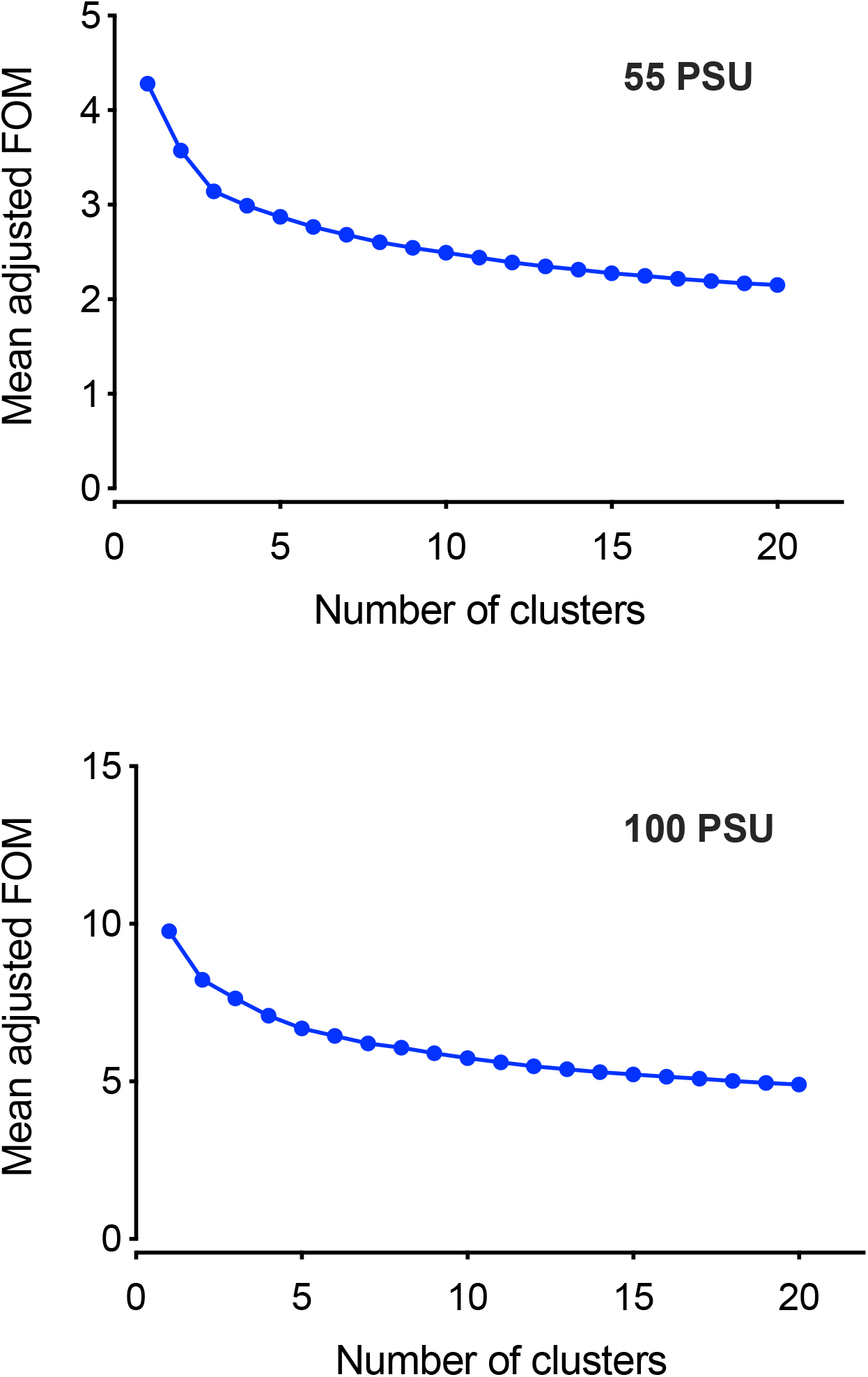
Figure of merit (FOM) analysis of temporal RNA-Seq dataset. The FOM analysis was performed for 20 iterations and the optimum number of clusters was found to be seven for both 55 and 100 PSU. This point was chosen for k-means clustering. Increasing the cluster size beyond the optimum number no longer lead information gain.

## References

An, B., Lan, J., Deng, X., Chen, S., Ouyang, C., Shi, H., Yang, J., Li, Y. 2017. Silencing of D-Lactate Dehydrogenase Impedes Glyoxalase System and Leads to Methylglyoxal Accumulation and Growth Inhibition in Rice. Frontiers in plant science, 8, 2071–2071.

Bartley, M.L., Boeing, W.J., Corcoran, A.A., Holguin, F.O., Schaub, T. 2013. Effects of salinity on growth and lipid accumulation of biofuel microalga *Nannochloropsis salina* and invading organisms. Biomass and Bioenergy, 54, 83–88.

Beck, J.G., Mathieu, D., Loudet, C., Buchoux, S., Dufourc, E.J. 2007. Plant sterols in “rafts”: a better way to regulate membrane thermal shocks. The FASEB Journal, 21(8), 1714–1723.

Bondioli, P., Della Bella, L., Rivolta, G., Zittelli, G.C., Bassi, N., Rodolfi, L., Casini, D., Prussi, M., Chiaramonti, D., Tredici, M.R. 2012. Oil production by the marine microalgae *Nannochloropsis* sp. F&M-M24 and Tetraselmis suecica F&M-M33. Bioresource technology, 114, 567–572.

Booth, W., Beardall, J. 1991. Effects of salinity on inorganic carbon utilization and carbonic anhydrase activity in the halotolerant alga *Dunaliella salina* (Chlorophyta). Phycologia, 30(2), 220–225.

Boyle, N.R., Page, M.D., Liu, B., Blaby, I.K., Casero, D., Kropat, J., Cokus, S.J., Hong-Hermesdorf, A., Shaw, J., Karpowicz, S.J. 2012. Three acyltransferases and nitrogen-responsive regulator are implicated in nitrogen starvation-induced triacylglycerol accumulation in *Chlamydomonas*. Journal of Biological Chemistry, 287(19), 15811–15825.

Brosché, M., Strid, Å. 1999. Cloning, Expression, and Molecular Characterization of a Small Pea Gene Family Regulated by Low Levels of Ultraviolet B Radiation and Other Stresses. Plant Physiology, 121(2), 479–488.

Černý, M., Kuklová, A., Hoehenwarter, W., Fragner, L., Novák, O., Rotková, G., Jedelský, P.L., Žáková, K., Šmehilová, M., Strnad, M., Weckwerth, W., Brzobohatý, B. 2013. Proteome and metabolome profiling of cytokinin action in *Arabidopsis* identifying both distinct and similar responses to cytokinin down- and up-regulation. Journal of Experimental Botany, 64(14), 4193–4206.

Chisti, Y. 2007. Biodiesel from microalgae. Biotechnology advances, 25(3), 294–306.

Choudhary, N., Sairam, R., Tyagi, A. 2005. Expression of Δ¹-pyrroline-5-carboxylate synthetase gene during drought in rice (*Oryza sativa* L.).

Coleman, J.R. 2000. Carbonic Anhydrase and Its Role in Photosynthesis. in: Photosynthesis: Physiology and Metabolism, (Eds.) R.C. Leegood, T.D. Sharkey, S. von Caemmerer, Springer Netherlands. Dordrecht, pp. 353–367.

Conde, A., Chaves, M.M., Gerós, H. 2011. Membrane Transport, Sensing and Signaling in Plant Adaptation to Environmental Stress. Plant and Cell Physiology, 52(9), 1583–1602.

Cornwall, C.E., Revill, A.T., Hurd, C.L. 2015. High prevalence of diffusive uptake of CO 2 by macroalgae in a temperate subtidal ecosystem. Photosynthesis Research, 124(2), 181–190.

Corteggiani Carpinelli, E., Telatin, A., Vitulo, N., Forcato, C., D’Angelo, M., Schiavon, R., Vezzi, A., Giacometti, G.M., Morosinotto, T., Valle, G. 2014. Chromosome scale genome assembly and transcriptome profiling of *Nannochloropsis gaditana* in nitrogen depletion. Mol Plant, 7(2), 323–35.

Courchesne, N.M.D., Parisien, A., Wang, B., Lan, C.Q. 2009. Enhancement of lipid production using biochemical, genetic and transcription factor engineering approaches. Journal of biotechnology, 141(1-2), 31–41.

Craigie, J., McLachlan, J. 1964. Glycerol as a photosynthetic product in *Dunaliella tertiolecta* Butcher. NATIONAL RESEARCH COUNCIL OF CANADA HALIFAX (NOVA SCOTIA) ATLANTIC REGIONAL LAB.

Demel, R.A., De Kruyff, B. 1976. The function of sterols in membranes. Biochimica et Biophysica Acta (BBA)-Reviews on Biomembranes, 457(2), 109–132.

Espie, G.S., Kimber, M.S. 2011. Carboxysomes: cyanobacterial RubisCO comes in small packages. Photosynth Res, 109(1-3), 7–20.

Ewels, P., Magnusson, M., Lundin, S., Käller, M. 2016. MultiQC: summarize analysis results for multiple tools and samples in a single report. Bioinformatics, 32(19), 3047–3048.

Fawley, M.W., Jameson, I., Fawley, K.P. 2015. The phylogeny of the genus *Nannochloropsis* (Monodopsidaceae, Eustigmatophyceae), with descriptions of *N. australis* sp. nov. and *Microchloropsis* gen. nov. Phycologia, 54(5), 545–552.

Fellenberg, K., Hauser, N.C., Brors, B., Neutzner, A., Hoheisel, J.D., Vingron, M. 2001. Correspondence analysis applied to microarray data. Proceedings of the National Academy of Sciences, 98(19), 10781–10786.

Flowers, T., Troke, P., Yeo, A. 1977. The mechanism of salt tolerance in halophytes. Annual review of plant physiology, 28(1), 89–121.

Flowers, T.J., Colmer, T.D. 2008. Salinity tolerance in halophytes. New Phytol, 179(4), 945–63.

Georgianna, D.R., Mayfield, S.P. 2012. Exploiting diversity and synthetic biology for the production of algal biofuels. Nature, 488(7411), 329.

Goyal, A. 2007. Osmoregulation in Dunaliella, Part II: Photosynthesis and starch contribute carbon for glycerol synthesis during a salt stress in *Dunaliella tertiolecta*. Plant Physiology and Biochemistry, 45(9), 705–710.

Green, P.E., Carmone, F.J., Smith, S.M. 1989. Multidimensional scaling: Concepts and applications. Allyn and Bacon.

Greenacre, M.J. 1984. Theory and applications of correspondence analysis.

Gu, N., Lin, Q., Li, G., Qin, G., Lin, J., Huang, L. 2012. Effect of salinity change on biomass and biochemical composition of *Nannochloropsis oculata*. Journal of the world aquaculture society, 43(1), 97–106.

Guarnieri, M.T., Nag, A., Smolinski, S.L., Darzins, A., Seibert, M., Pienkos, P.T. 2011. Examination of triacylglycerol biosynthetic pathways via de novo transcriptomic and proteomic analyses in an unsequenced microalga. PLoS One, 6(10), e25851.

Helbling, E.W., Buma, A.G., Boelen, P., Van der Strate, H.J., Giordanino, M.V.F., Villafañe, V.E. 2011. Increase in Rubisco activity and gene expression due to elevated temperature partially counteracts ultraviolet radiation–induced photoinhibition in the marine diatom *Thalassiosira weissflogii*. Limnology and Oceanography, 56(4), 1330–1342.

Hibino, T., Waditee, R., Araki, E., Ishikawa, H., Aoki, K., Tanaka, Y., Takabe, T. 2002. Functional characterization of choline monooxygenase, an enzyme for betaine synthesis in plants. Journal of Biological Chemistry, 277(44), 41352–41360.

Ho, S.-H., Nakanishi, A., Kato, Y., Yamasaki, H., Chang, J.-S., Misawa, N., Hirose, Y., Minagawa, J., Hasunuma, T., Kondo, A. 2017. Dynamic metabolic profiling together with transcription analysis reveals salinity-induced starch-to-lipid biosynthesis in alga *Chlamydomonas* sp. JSC4. Scientific reports, 7, 45471.

Hu, C., Delauney, A.J., Verma, D. 1992. A bifunctional enzyme (delta 1-pyrroline-5-carboxylate synthetase) catalyzes the first two steps in proline biosynthesis in plants. Proceedings of the National Academy of Sciences, 89(19), 9354–9358.

Huertas, E., Montero, O., Lubián, L.M. 2000. Effects of dissolved inorganic carbon availability on growth, nutrient uptake and chlorophyll fluorescence of two species of marine microalgae. Aquacultural engineering, 22(3), 181–197.

Ihnken, S., Eggert, A., Beardall, J. 2010. Exposure times in rapid light curves affect photosynthetic parameters in algae. Aquatic Botany, 93(3), 185–194.

Jessen, D., Roth, C., Wiermer, M., Fulda, M. 2015. Two activities of long-chain acyl-coenzyme A synthetase are involved in lipid trafficking between the endoplasmic reticulum and the plastid in *Arabidopsis*. Plant physiology, 167(2), 351–366.

Jing, H., Takagi, J., Liu, J.-h., Lindgren, S., Zhang, R.-g., Joachimiak, A., Wang, J.-h., Springer, T.A. 2002. Archaeal Surface Layer Proteins Contain β Propeller, PKD, and β Helix Domains and Are Related to Metazoan Cell Surface Proteins. Structure, 10(10), 1453–1464.

Johnson, X., Alric, J. 2013. Central carbon metabolism and electron transport in *Chlamydomonas reinhardtii*: metabolic constraints for carbon partitioning between oil and starch. Eukaryotic cell, 12(6), 776–793.

Kachroo, A., Kachroo, P. 2009. Fatty Acid-derived signals in plant defense. Annu Rev Phytopathol, 47, 153–76.

Karthikaichamy, A., Beardall, J., Coppel, R., Noronha, S., Bulach, D., Schittenhelm, R.B. and Srivastava, S., 2020. A Data-Independent-Acquisition-based proteomic approach towards understanding the acclimation strategy of Microchloropsis gaditana CCMP526 in hypersaline conditions. bioRxiv.

Karthikaichamy, A., Deore, P., Srivastava, S., Coppel, R., Bulach, D., Beardall, J., Noronha, S. 2018. Temporal acclimation of *Microchloropsis gaditana* CCMP526 in response to hypersalinity. Bioresource Technology, 254, 23–30.

Kemble, A., Macpherson, H.T. 1954. Liberation of amino acids in perennial rye grass during wilting. Biochemical Journal, 58(1), 46.

Kim, B.-H., Ramanan, R., Kang, Z., Cho, D.-H., Oh, H.-M., Kim, H.-S. 2016. *Chlorella sorokiniana* HS1, a novel freshwater green algal strain, grows and hyperaccumulates lipid droplets in seawater salinity. Biomass and Bioenergy, 85, 300–305.

Kishino, H., Waddell, P.J. 2000. Correspondence analysis of genes and tissue types and finding genetic links from microarray data. Genome Informatics, 11, 83–95.

Kravchik, M., Bernstein, N. 2013. Effects of salinity on the transcriptome of growing maize leaf cells point at cell-age specificity in the involvement of the antioxidative response in cell growth restriction. BMC Genomics, 14, 24.

Li, X., Benning, C., Kuo, M.-H. 2012a. Rapid triacylglycerol turnover in *Chlamydomonas reinhardtii* requires a lipase with broad substrate specificity. Eukaryotic cell, EC. 00268-12.

Li, X., Moellering, E.R., Liu, B., Johnny, C., Fedewa, M., Sears, B.B., Kuo, M.-H., Benning, C. 2012b. A galactoglycerolipid lipase is required for triacylglycerol accumulation and survival following nitrogen deprivation in *Chlamydomonas reinhardtii*. The Plant Cell, tpc. 112.105106.

Liu, J., Han, D., Yoon, K., Hu, Q., Li, Y. 2016. Characterization of type 2 diacylglycerol acyltransferases in *Chlamydomonas reinhardtii* reveals their distinct substrate specificities and functions in triacylglycerol biosynthesis. The Plant Journal, 86(1), 3–19.

Madsen, T., Baattrup-Pedersen, A. 1995. Regulation of growth and photosynthetic performance in *Elodea canadensis* in response to inorganic nitrogen. Functional Ecology, 239–247.

McLean, M.D., Yevtushenko, D.P., Deschene, A., Van Cauwenberghe, O.R., Makhmoudova, A., Potter, J.W., Bown, A.W., Shelp, B.J. 2003. Overexpression of glutamate decarboxylase in transgenic tobacco plants confers resistance to the northern root-knot nematode. Molecular Breeding, 11(4), 277–285.

Mi, H., Huang, X., Muruganujan, A., Tang, H., Mills, C., Kang, D., Thomas, P.D. 2017. PANTHER version 11: expanded annotation data from Gene Ontology and Reactome pathways, and data analysis tool enhancements. Nucleic Acids Research, 45(D1), D183–D189.

Molnár, I., Lopez, D., Wisecaver, J.H., Devarenne, T.P., Weiss, T.L., Pellegrini, M., Hackett, J.D. 2012. Bio-crude transcriptomics: gene discovery and metabolic network reconstruction for the biosynthesis of the terpenome of the hydrocarbon oil-producing green alga, *Botryococcus braunii* race B (Showa). BMC genomics, 13(1), 576.

Moody, J.W., McGinty, C.M., Quinn, J.C. 2014. Global evaluation of biofuel potential from microalgae. Proceedings of the National Academy of Sciences, 111(23), 8691–8696.

Olvera-Carrillo, Y., Luis Reyes, J., Covarrubias, A.A. 2011. Late embryogenesis abundant proteins: versatile players in the plant adaptation to water limiting environments. Plant signaling & behavior, 6(4), 586–589.

Omidbakhshfard, M.A., Omranian, N., Ahmadi, F.S., Nikoloski, Z., Mueller-Roeber, B. 2012. Effect of salt stress on genes encoding translation-associated proteins in *Arabidopsis thaliana*. Plant Signaling & Behavior, 7(9), 1095–1102.

Ou, X., Ji, C., Han, X., Zhao, X., Li, X., Mao, Y., Wong, L.L., Bartlam, M., Rao, Z. 2006. Crystal structures of human glycerol 3-phosphate dehydrogenase 1 (GPD1). J Mol Biol, 357(3), 858–69.

Pandit, P.R., Fulekar, M.H., Karuna, M.S.L. 2017. Effect of salinity stress on growth, lipid productivity, fatty acid composition, and biodiesel properties in Acutodesmus obliquus and *Chlorella vulgaris*. Environmental Science and Pollution Research, 24(15), 13437–13451.

Perrineau, M.M., Zelzion, E., Gross, J., Price, D.C., Boyd, J., Bhattacharya, D. 2014. Evolution of salt tolerance in a laboratory reared population of *Chlamydomonas reinhardtii*. Environmental microbiology, 16(6), 1755–1766.

Pical, C., Westergren, T., Dove, S.K., Larsson, C., Sommarin, M. 1999. Salinity and hyperosmotic stress induce rapid increases in phosphatidylinositol 4, 5-bisphosphate, diacylglycerol pyrophosphate, and phosphatidylcholine in *Arabidopsis thaliana* cells. Journal of Biological Chemistry, 274(53), 38232–38240.

Posé, D., Castanedo, I., Borsani, O., Nieto, B., Rosado, A., Taconnat, L., Ferrer, A., Dolan, L., Valpuesta, V., Botella, M.A. 2009. Identification of the *Arabidopsis* dry2/sqe1-5 mutant reveals a central role for sterols in drought tolerance and regulation of reactive oxygen species. The Plant Journal, 59(1), 63–76.

Powell, D.R., 2015. Degust v3. 2.0. Zenodo https://doi.org/10.5281/zenodo, 3258933.

Reed, M., Graham, D. 1981. Carbonic anhydrase in plants: distribution, properties and possible physiological roles. Progress in phytochemistry.

Rismani-Yazdi, H., Haznedaroglu, B.Z., Bibby, K., Peccia, J. 2011. Transcriptome sequencing and annotation of the microalgae *Dunaliella tertiolecta*: pathway description and gene discovery for production of next-generation biofuels. BMC genomics, 12(1), 148.

Rismani-Yazdi, H., Haznedaroglu, B.Z., Hsin, C., Peccia, J. 2012. Transcriptomic analysis of the oleaginous microalga *Neochloris oleoabundans* reveals metabolic insights into triacylglyceride accumulation. Biotechnology for Biofuels, 5(1), 74.

Rossmann, M., Liljas, A., Brändén, C., Banaszak, L. 1975. The Enzymes, edited by PD Boyer, Vol. XI, New York: Academic Press.

Saeed, A., Sharov, V., White, J., Li, J., Liang, W., Bhagabati, N., Braisted, J., Klapa, M., Currier, T., Thiagarajan, M. 2003. TM4: a free, open-source system for microarray data management and analysis. Biotechniques, 34(2), 374–378.

Saradhi, P.P., AliaArora, S., Prasad, K. 1995. Proline accumulates in plants exposed to UV radiation and protects them against UV-induced peroxidation. Biochemical and biophysical research communications, 209(1), 1–5.

Schaller, H. 2010. Sterol and steroid biosynthesis and metabolism in plants and microorganisms.

Schat, H., Sharma, S.S., Vooijs, R. 1997. Heavy metal-induced accumulation of free proline in a metal-tolerant and a nontolerant ecotype of *Silene vulgaris*. Physiologia Plantarum, 101(3), 477–482.

Scholz, M.J., Weiss, T.L., Jinkerson, R.E., Jing, J., Roth, R., Goodenough, U., Posewitz, M.C., Gerken, H.G. 2014. Ultrastructure and Composition of the *Nannochloropsis gaditana* Cell Wall. Eukaryotic Cell, 13(11), 1450–1464.

Siaut, M., Cuine, S., Cagnon, C., Fessler, B., Nguyen, M., Carrier, P., Beyly, A., Beisson, F., Triantaphylides, C., Li-Beisson, Y. 2011. Oil accumulation in the model green alga *Chlamydomonas reinhardtii*: characterization, variability between common laboratory strains and relationship with starch reserves. BMC biotechnology, 11(1), 7.

Silvestro, D., Andersen, T.G., Schaller, H., Jensen, P.E. 2013. Plant sterol metabolism. Δ7-sterol-C5-desaturase (STE1/DWARF7), Δ5, 7-sterol-Δ7-reductase (DWARF5) and Δ24-sterol-Δ24-reductase (DIMINUTO/DWARF1) show multiple subcellular localizations in *Arabidopsis thaliana* (Heynh) L. PloS one, 8(2), e56429.

Simionato, D., Block, M.A., La Rocca, N., Jouhet, J., Maréchal, E., Finazzi, G., Morosinotto, T. 2013. The response of *Nannochloropsis gaditana* to nitrogen starvation includes de novo biosynthesis of triacylglycerols, a decrease of chloroplast galactolipids, and reorganization of the photosynthetic apparatus. Eukaryotic cell, 12(5), 665–676.

Simionato, D., Sforza, E., Carpinelli, E.C., Bertucco, A., Giacometti, G.M., Morosinotto, T. 2011. Acclimation of *Nannochloropsis gaditana* to different illumination regimes: effects on lipids accumulation. Bioresource technology, 102(10), 6026–6032.

Siripornadulsil, S., Traina, S., Verma, D.P.S., Sayre, R.T. 2002. Molecular mechanisms of proline-mediated tolerance to toxic heavy metals in transgenic microalgae. The Plant Cell, 14(11), 2837–2847.

Sonawane, P.D., Heinig, U., Panda, S., Gilboa, N.S., Yona, M., Kumar, S.P., Alkan, N., Unger, T., Bocobza, S., Pliner, M., Malitsky, S., Tkachev, M., Meir, S., Rogachev, I., Aharoni, A. 2018. Short-chain dehydrogenase/reductase governs steroidal specialized metabolites structural diversity and toxicity in the genus *Solanum*. Proceedings of the National Academy of Sciences, 115(23), E5419–E5428.

Spreitzer, R.J. 2003. Role of the small subunit in ribulose-1,5-bisphosphate carboxylase/oxygenase. Arch Biochem Biophys, 414(2), 141–9.

Stephens, E., Ross, I.L., King, Z., Mussgnug, J.H., Kruse, O., Posten, C., Borowitzka, M.A., Hankamer, B. 2010. An economic and technical evaluation of microalgal biofuels. Nature biotechnology, 28(2), 126.

Sukenik, A. 1991. Ecophysiological considerations in the optimization of eicosapentaenoic acid production by *Nannochloropsis* sp.(Eustigmatophyceae). Bioresource Technology, 35(3), 263–269.

Sukenik, A., Carmeli, Y., Berner, T. 1989. Regulation of fatty acid composition by irradiance level in the eustigmatophyte *Nannochloropsis* sp. 1. Journal of Phycology, 25(4), 686–692.

Szabados, L., Kovács, H., Zilberstein, A., Bouchereau, A. 2011. Plants in extreme environments: importance of protective compounds in stress tolerance. in: Advances in Botanical Research, Vol. 57, Elsevier, pp. 105–150.

Takabe, T., Incharoensakdi, A., Arakawa, K., Yokota, S. 1988. CO2 fixation rate and RuBisCO content increase in the halotolerant cyanobacterium, *Aphanothece halophytica*, grown in high salinities. Plant physiology, 88(4), 1120–1124.

Takagi, M., Yoshida, T. 2006. Effect of salt concentration on intracellular accumulation of lipids and triacylglyceride in marine microalgae *Dunaliella* cells. Journal of bioscience and bioengineering, 101(3), 223–226.

Tortell, P.D., Martin, C.L., Corkum, M.E. 2006. Inorganic carbon uptake and intracellular assimilation by subarctic Pacific phytoplankton assemblages. Limnology and oceanography, 51(5), 2102–2110.

Tsyganov, K., Pery, A.J., Archer, S.K. and Powel, D., 2018. RNAsik: A Pipeline for complete and reproducible RNA-seq analysis that runs anywhere with speed and ease. Journal of Open Source Software, 3(28), p.583.

Verbruggen, N., Villarroel, R., Van Montagu, M. 1993. Osmoregulation of a pyrroline-5-carboxylate reductase gene in *Arabidopsis thaliana*. Plant Physiology, 103(3), 771–781.

Wada, H., Gombos, Z., Murata, N. 1994. Contribution of membrane lipids to the ability of the photosynthetic machinery to tolerate temperature stress. Proc Natl Acad Sci U S A, 91(10), 4273–7.

Wasylenko, T.M., Ahn, W.S., Stephanopoulos, G. 2015. The oxidative pentose phosphate pathway is the primary source of NADPH for lipid overproduction from glucose in *Yarrowia lipolytica*. Metab Eng, 30, 27–39.

Wegmann, K. 1971. Osmotic regulation of photosynthetic glycerol production in *Dunaliella*. Biochimica et Biophysica Acta (BBA)-Bioenergetics, 234(3), 317–323.

Wei, J., Ji, H., Guo, M., Qin, Q. 2012. Isolation and characterization of a thioredoxin domain-containing protein 12 from orange-spotted grouper, *Epinephelus coioides*. Fish Shellfish Immunol, 33(3), 667–73.

Wijffels, R.H., Barbosa, M.J. 2010. An outlook on microalgal biofuels. Science, 329(5993), 796–799.

Wyn Jones, R. 1977. A hypothesis on cytoplasmic osmoregulation. Regulation of Cell Membrane Activities in Higher Plants., 121–136.

Xue, J., Wang, L., Zhang, L., Balamurugan, S., Li, D.-W., Zeng, H., Yang, W.-D., Liu, J.-S., Li, H.-Y. 2016. The pivotal role of malic enzyme in enhancing oil accumulation in green microalga *Chlorella pyrenoidosa*. Microbial cell factories, 15(1), 120.

Yamada, M., Morishita, H., Urano, K., Shiozaki, N., Yamaguchi-Shinozaki, K., Shinozaki, K., Yoshiba, Y. 2005. Effects of free proline accumulation in petunias under drought stress. Journal of Experimental Botany, 56(417), 1975–1981.

Yang, F., Xiang, W., Li, T., Long, L. 2018. Transcriptome analysis for phosphorus starvation-induced lipid accumulation in *Scenedesmus* sp. Scientific reports, 8.

Yeagle, P. 1991. Modulation of membrane function by cholesterol. Biochimie, 73(10), 1303–1310.

Yoshiba, Y., Kiyosue, T., Katagiri, T., Ueda, H., Mizoguchi, T., Yamaguchi-Shinozaki, K., Wada, K., Harada, Y., Shinozaki, K. 1995. Correlation between the induction of a gene for delta 1-pyrroline-5-carboxylate synthetase and the accumulation of proline in *Arabidopsis thaliana* under osmotic stress. Plant J, 7(5), 751–60.

Young, J.N., Hopkinson, B.M. 2017. The potential for co-evolution of CO2-concentrating mechanisms and Rubisco in diatoms. Journal of experimental botany, 68(14), 3751–3762.

Yu, S., Zhang, X., Guan, Q., Takano, T., Liu, S. 2007. Expression of a carbonic anhydrase gene is induced by environmental stresses in rice (*Oryza sativa* L.). Biotechnol Lett, 29(1), 89–94.

Zeng, X., Jin, P., Xia, J., Liu, Y. 2019. Correlation of carbonic anhydrase and Rubisco in the growth and photosynthesis in the diatom *Phaeodactylum tricornutum*. Journal of Applied Phycology, 31(1), 123–129.

Zhang, Y., Adams, I.P., Ratledge, C. 2007. Malic enzyme: the controlling activity for lipid production? Overexpression of malic enzyme in *Mucor circinelloides* leads to a 2.5-fold increase in lipid accumulation. Microbiology, 153(7), 2013–2025.

Zhu, B.-H., Zhang, R.-H., Lv, N.-N., Yang, G.-P., Wang, Y.-S., Pan, K.-H. 2018. The Role of Malic Enzyme on Promoting Total Lipid and Fatty Acid Production in *Phaeodactylum tricornutum*. Frontiers in Plant Science, 9(826).

Zörb, C., Schmitt, S., Neeb, A., Karl, S., Linder, M., Schubert, S. 2004. The biochemical reaction of maize (*Zea mays* L.) to salt stress is characterized by a mitigation of symptoms and not by a specific adaptation. Plant Science, 167(1), 91–100.

